# Label free multimodal optical imaging of metabolic heterogeneity and correlation in aging by integrating SRS, MPF, FLIM, and SHG

**DOI:** 10.64898/2026.04.11.717872

**Authors:** Hongje Jang, Shuang Wu, Hyehyun Kim, Tong-You Wade Wei, Jian Kang, Anahita Ataran, Fangyuan Gao, Dorota Skowronska-Krawczyk, Kenneth B. Margulies, Ali Javaheri, Wei Sun, John Y-J Shyy, Erkin Seker, Lingyan Shi

## Abstract

Cellular metabolism is governed by the coordinated organization of macromolecules, including lipids and proteins, together with redox-active cofactors such as NADH and FAD. However, resolving these biochemical features quantitatively and spatially at subcellular resolution remains challenging because no single imaging modality can capture molecular composition, redox state, and tissue architecture simultaneously without labeling. Here, we present **MANIFEST** (**M**ultimodal **A**nalysis of **N**onlinear **I**maging for **F**unctional **E**ndogenous **S**patial correla**T**ion), a label-free imaging platform that integrates stimulated Raman scattering (SRS), second harmonic generation (SHG), multiphoton fluorescence (MPF), and fluorescence lifetime imaging microscopy (FLIM) with their spatial correlation. MANIFEST combines chemical imaging of lipids with autofluorescence- and lifetime-based quantification of NADH and FAD metabolism, enabling spatially resolved analysis of metabolic heterogeneity at organelle and tissue-compartment levels. We apply this framework to four distinct aging or disease models: amyloid-beta-treated tri-cultured brain cells, high-fat diet mouse liver, human non-ischemic cardiomyopathy tissue, and aging mouse retina. Across these systems, MANIFEST reveals disease-associated lipid remodeling, redox imbalance, disrupted metabolic zonation, collagen reorganization, and layer-specific metabolic changes. By integrating complementary nonlinear optical modalities into a single label-free platform, MANIFEST provides an extensible, proof-of-concept platform for high-resolution metabolic phenotyping in complex biological systems and offers new opportunities for studying disease mechanisms, aging biology, and metabolism-driven tissue pathology.

## 1. Introduction

Metabolic heterogeneity is a defining feature of both normal physiology and disease progression. Across aging tissues and pathological states, cells differ in their lipid composition, protein organization, redox balance, and extracellular matrix remodeling. These differences often emerge at subcellular, cellular, and tissue-compartment scales. Such biochemical variation is closely linked to functional state and disease trajectory in diverse systems, including neurodegeneration, metabolic disease, cardiovascular dysfunction, and retinal aging. However, directly visualizing these metabolic features in situ remains challenging because no single imaging modality can simultaneously resolve molecular composition, cofactor dynamics, and tissue structure with high spatial resolution and without exogenous labeling[1].

Current imaging strategies each capture only part of this landscape. Conventional fluorescence microscopy offers high sensitivity and molecular specificity[2], but it often depends on exogenous labels, is constrained in multiplexing capacity[3], and may perturb the biological system under study. Super-resolution fluorescence imaging methods can achieve exceptional spatial precision[4–11], yet this gain in specificity often comes at the expense of throughput, molecular breadth, and compatibility with large-scale tissue analysis. By contrast, label-free optical approaches offer the possibility of probing endogenous biochemical information directly, but most single-modality methods remain limited in the range of molecular features they can report. As a result, there is a substantial unmet need for imaging platforms that can quantitatively integrate multiple endogenous contrasts and map them across intact biological samples in a unified framework.

Nonlinear optical microscopy provides a powerful foundation for such integration because distinct light–matter interactions generate complementary biochemical readouts[12–14]. Stimulated Raman scattering (SRS) microscopy enables label-free imaging of major biomolecular classes, particularly lipids and proteins. Hyperspectral SRS (HS-SRS) can further distinguish molecular subtypes through vibrational spectral signatures[15–18]. Second harmonic generation (SHG) selectively visualizes non-centrosymmetric structures such as collagen[19, 20], providing structural information on extracellular matrix organization. Multiphoton fluorescence (MPF) detects endogenous fluorophores, including the metabolic cofactors NADH and FAD, while fluorescence lifetime imaging microscopy (FLIM) resolves their microenvironment-dependent lifetime states, enabling more detailed interpretation of redox and metabolic activity[21–23]. Together, these modalities are highly complementary: SRS reports chemical composition, SHG captures structural organization, and MPF/FLIM provide functional readouts of metabolic cofactors. Integrating them into a single analytical framework can therefore reveal relationships between macromolecular composition and metabolic states that are inaccessible to any one modality alone.

Recent advances in computational imaging have further expanded the utility of nonlinear optical methods. HS-SRS analysis has enabled quantitative mapping of lipid subtypes, while image processing approaches have improved the spatial resolution of SRS and MPF[18, 24–26]. Nevertheless, most of these developments have emphasized macromolecular composition, whereas small metabolic cofactors remain under-characterized in relation to surrounding lipids, proteins, and tissue architecture. This limitation is especially important because NADH and FAD are central reporters of cellular metabolism. In particular, NADH exists in multiple functional states, including free NADH, enzyme-bound NADH, and NADPH-associated pools, each reflecting distinct biochemical activities[27–30]. A multimodal strategy that quantitatively links these cofactor states to endogenous molecular distributions would therefore provide a more complete view of metabolic heterogeneity in cells and tissues.

Despite the strong potential of multimodal imaging approaches and advanced computational imaging techniques for quantifying metabolic activities, their combined application to metabolism studies has yet to be established. In the context of medical diagnostics, multimodal imaging approaches have been employed to identify pathological regions within tissues based on compositional contrast, without the need for exogenous labeling[31–33]. With a properly designed measurement framework and suitable analytical methods, multimodal imaging can enable the simultaneous characterization of multiple metabolic activities. Here, we present **MANIFEST** (**M**ultimodal **A**nalysis of **N**onlinear **I**maging for **F**unctional **E**ndogenous **S**patial correla**T**ion), a label-free multimodal imaging platform that integrates SRS, SHG, MPF, and FLIM for spatially resolved metabolic profiling. MANIFEST combines SRS imaging of lipid distributions with autofluorescence- and lifetime-based analysis of NADH and FAD, enabling quantitative assessment of both macromolecular organization and cofactor-associated metabolic state (anabolic versus catabolic states) within the same biological specimen. By incorporating phasor analysis together with fluorescence lifetime curve analysis, the platform resolves detailed information from endogenous fluorophores and relates these measurements to lipid-rich organelles and tissue compartments. At the end of the MANIFEST workflow, by mapping the spatial correlation of each metabolic activity and biochemical distribution, we can identify the correlated metabolic activities and competing activities. We apply MANIFEST across four distinct models of disease or aging: amyloid-beta-treated tri-cultured brain cells, high-fat diet mouse liver, human non-ischemic cardiomyopathy tissue, and aging mouse retina. Across these systems, MANIFEST reveals spatially localized lipid remodeling, redox imbalance, altered collagen organization, disrupted metabolic zonation, and tissue-specific structural reorganization. These proof-of-concept results establish MANIFEST as an extensible framework for high-resolution, label-free analysis of metabolic heterogeneity in complex biological systems.

## 2. Materials and Methods

### 2.1 Imaging Setup for MANIFEST

MANIFEST was implemented on a custom-built upright laser-scanning microscope (Olympus) configured for stimulated Raman scattering (SRS), second harmonic generation (SHG), multiphoton fluorescence (MPF), and fluorescence lifetime imaging microscopy (FLIM). A 25× water-immersion objective lens (XLPLN WMP2, 1.05 NA, Olympus) was used to maximize near-infrared throughput. Excitation was provided by a picoEmerald system (Applied Physics & Electronics) delivering a synchronized pulsed pump beam tunable from 720 to 990 nm (5–6 ps pulse width, 80 MHz repetition rate) together with a 1032 nm Stokes beam (6 ps pulse width, 80 MHz repetition rate). The pump and Stokes beams were coupled into the microscope and focused onto the sample. Transmitted signals were collected by a high-NA oil condenser (1.4 NA). To isolate the stimulated Raman loss signal, a high-optical-density 950 nm short-pass filter (Thorlabs) was used to block the Stokes beam completely while transmitting the pump beam to a silicon photodiode. The output current from the photodiode was terminated, filtered, and demodulated at 20 MHz using a lock-in amplifier. The demodulated signal was then reconstructed into images using the FV-OSR module in FV3000 software (Olympus)[1, 18, 34-36]. Cardiac tissue and cell images were acquired at 512 × 512 pixels, whereas liver and retinal tissue images were acquired at 1024 × 1024 pixels. The pixel dwell time was 20 μs. MPF, SHG, and FLIM measurements were integrated into the same microscope and acquired from the same regions of interest using a photomultiplier tube (PMT) and a time-correlated single-photon counting (TCSPC) module for FLIM acquisition.

### 2.2 Correlation Coefficient Mapping

As described in Fig. 1d, PCC in each pixel is calculated. The equation to calculate PCC from two images of A and B is as follows.

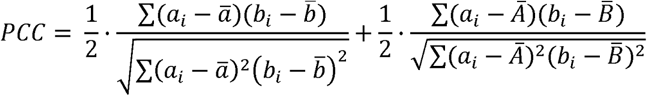

**Fig. 1.**
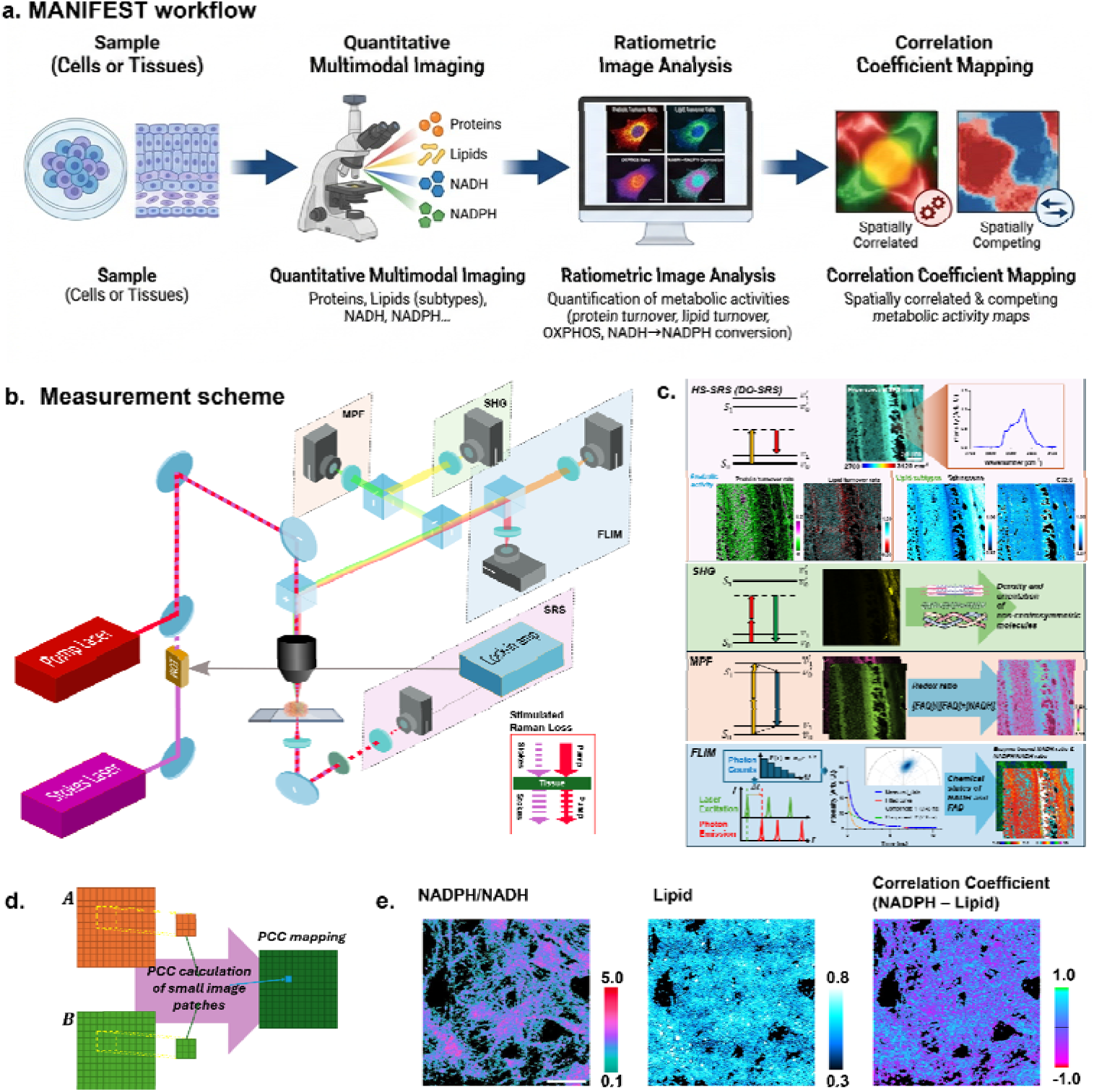
MANIFEST: multimodal nonlinear imaging of metabolic heterogeneity. **a.** Workflow of MANIFEST. Using non-linear multimodal imaging approach, images representing various molecular spatial distribution are measured. From the images, multiple metabolic activity distribution are shown as a form of ratiometric images. By generating correlation coefficient maps of the metabolic activities, correlations of the metabolic activities are studied. **b.** Schematic of the multimodal imaging platform of MANIFEST, which integrates stimulated Raman scattering (SRS), second harmonic generation (SHG), multiphoton fluorescence (MPF), and fluorescence lifetime imaging microscopy (FLIM) on a single microscope framework. Pump and Stokes beams are delivered to the sample in a confocal configuration. SRS signals are detected in transmission, whereas back-propagating photons are separated by chromatic beam splitters and directed to dedicated detectors for SHG, MPF, and FLIM. SHG is generated using the Stokes beam while minimizing two-photon excitation of NADH, whereas the pump beam enables two-photon excitation of both NADH and FAD. Images from all modalities were acquired sequentially from the same region of interest. **c.** Analytical workflow of MANIFEST. Hyperspectral SRS images acquired in the C–H and C–D spectral regions were used to quantify lipid composition, lipid subtype distribution, and relative protein and lipid turnover. SHG images report the spatial distribution of non-centrosymmetric structures, including collagen fibers. MPF images of NADH and FAD were used to calculate the optical redox ratio. FLIM data were analyzed by phasor analysis followed by biexponential fitting to resolve distinct fluorescence lifetime states of NADH and FAD. **d.** Schematic of spatial correlation mapping method. From the small image patches of two images, Pearson’s correlation coefficient (PCC) is calculated, and the PCC is stored in each representing pixel. By repeating this process pixel-by-pixel, correlation coefficient map is generated. **e.** Example of PCC map between NADPH ratio and lipid amount. The positively correlated ROIs are expressed as blue and cyan colors, and negatively correlated ROIs are represented as red and magenta colors.

*α_i_* and *b_i_* are intensities in each pixel *ā* and *b̄* are averaged intensities of each image patches. *Ā* and *B̄* are averaged intensities of whole ROIs of each image. To balance the local and global intensity change information, the two different PCCs were averaged.

### 2.3 FLIM Analysis

The FLIM data were processed using custom-built Python scripts for phasor plot[23] and lifetime curve fitting. As described in Fig. S1, a two-step analysis approach was employed. First, an uncalibrated phasor diagram was generated. From this phasor diagram, pixels representing different chemical environments were clustered using the Mean Shift algorithm[37]. Following clustering, the photon-counting histogram of each pixel cluster was averaged and fitted with a biexponential decay model. By using these fitting results as initial values for the precise fitting of individual pixel lifetime curves, a detailed two-component analysis of free and enzyme-bound flavin and NADH was achieved. Phasor positions were used only for clustering and for initializing the fits and were not interpreted quantitatively; lifetime values were derived exclusively from the biexponential fits. Through the measurement of FluoSpheres Polystyrene Microspheres (F13081, Thermo Fisher, UK), the instrument response function was accounted[38]. After generating the binary filter from MPF image, the pixels have meaningful intensity were selected for the analysis. To keep the quality of bi-exponential fitting, Finally, the lifetime of enzyme-bound NADH was used to calculate the NADPH/NADH ratio[39].

### 2.4 Image preprocessing and LD analysis

All acquired images, except for the FLIM data, were denoised using the PURE-Denoise filter[40, 41]. Following denoising, the water signal from the hyperspectral SRS images was removed according to the method detailed in Fig. S2. The signal intensities in each image channel were then analyzed. For LD analysis, individual droplets were quantified using the 3D Objects Counter plugin[42] in ImageJ. Based on parameters including position, volume, surface area, and mean distance, LD masks were generated. Using these masks, the signals from each channel were averaged and plotted in Figs. 2 and 4.

**Fig. 2.**
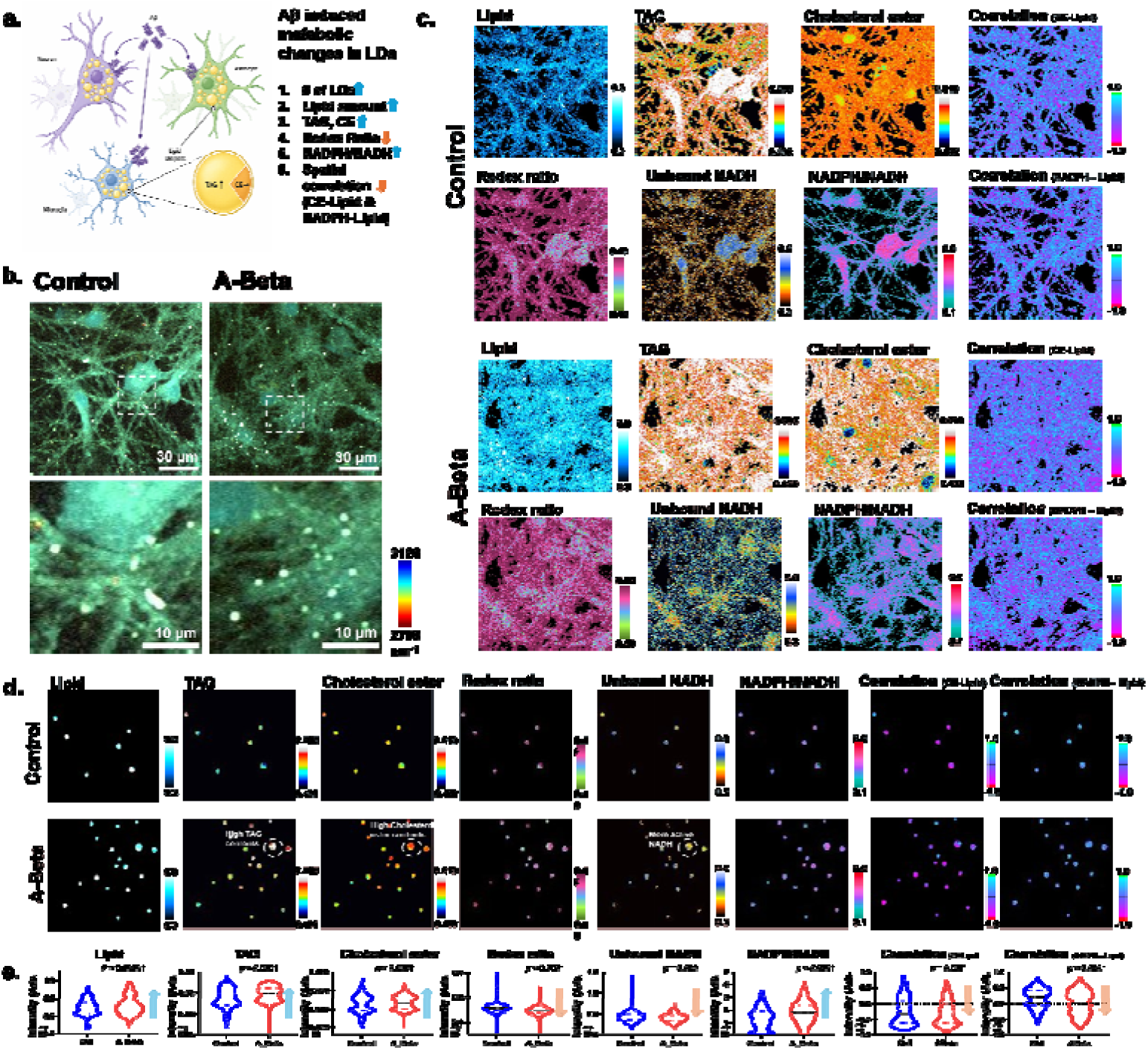
Amyloid-beta treatment remodels lipid-droplet composition and cofactor-associated metabolism in tri-cultured brain cells. **a.** Schematic summarizes the amyloid-beta (Aβ) induced metabolic changes in lipid droplets (LDs). **b.** Hyperspectral SRS images show the distribution of cells in each field of view. Enlarged views of the boxed regions, showing heterogeneity among individual lipid droplets (LDs). **c.** Multimodal images of control and Aβ-treated tri-cultures containing neurons, astrocytes, and microglia. From hyperspectral SRS images, ratiometric distributions of triacylglycerol (TAG) and cholesterol ester were generated. MPF and FLIM images report the optical redox ratio, unbound NADH ratio, and NADPH-associated signal. Images were acquired from three independent culture preparations per group, with five regions of interest (ROIs) imaged per culture preparation. ROIs from the same culture preparation were treated as technical sampling units. **d.** Representative droplets exhibiting the dominant compositional and metabolic trends are marked with red circles. **e.** Quantification of LD-associated signals in control and Aβ-treated cultures. Compared with control cultures, LDs in Aβ-treated cells showed higher TAG, cholesterol ester, and total lipid signals, together with altered redox ratio, reduced unbound NADH ratio, and elevated NADPH-associated signal. The spatial correlations of cholesterol ester – lipid amount and NADPH ratio – lipid amount were decreased to more negative value. For group comparisons, measurements from multiple ROIs within the same culture preparation were averaged to generate one value per biological replicate before statistical analysis (n = 3 biological replicates per condition; 15 ROIs in total per condition). LD counts are shown for descriptive distributional analysis only and were not treated as independent biological replicates (571 LDs in control; 901 LDs in Aβ-treated cultures).

### 2.5 Image analysis for DO-SRS and PRM-SRS

Lipid subtype composition was determined using the PRM-SRS method[18]. Hyperspectral images covering the C-H vibration region (2700-3120 cm^-1^) were acquired at 6 cm^-1^ intervals, yielding a stack of 71 frames per field of view. Spectral similarities between the acquired data and a reference library, identical to that used in the previous PRM-SRS study, were calculated to identify the lipid subtypes present in each sample. The results were expressed as normalized ratiometric images for each lipid subtype, computed as [target lipid] / ([target lipid] + [PE]).

Hyperspectral images in the C-D vibration region (2000–2300 cm^-1^) were acquired at 6 cm^-1^ intervals, producing 51 frames per stack. From these stacks, newly synthesized lipid images (2135 cm^-1^) and background images (2000 cm^-1^) were extracted. To correct baseline intensity offsets, the background images were subtracted from the newly synthesized lipid images. The corrected images were then divided by the existing lipid channel image, derived from the C-H hyperspectral stack, to convert absolute new-lipid signals into lipid turnover rate maps. Because a single labeling duration was used and no kinetic model was fitted, these maps report relative D-label incorporation (normalized new-lipid signal) rather than absolute kinetic turnover rates[18], and the term “lipid turnover” is used in this relative sense throughout.

### 2.6 Primary rat cortical tri-cultures and Aβ treatment

Primary cortical cells were isolated from postnatal day 0-2 Sprague-Dawley rat pups and maintained as previously described[43]. Briefly, cells were plated on poly-L-lysine-coated surfaces and maintained in Neurobasal-A-based plus media supplemented with B27 and GlutaMAX. A neuron-glia co-culture medium was used as the base, and triculture conditions were established by supplementing the medium with IL-34, TGF-β2, and cholesterol to support microglial survival, following published protocols[43]. FITC-tagged Aβ42 was prepared as previously reported[43], aliquoted, and stored at −80 °C. Prior to treatment, Aβ was incubated overnight at 4 °C to allow aggregation. At DIV −10, cultures were treated by replacing half of the medium with Aβ-containing medium at twice the final concentration, while vehicle controls received equivalent DMSO in DPBS+. Cultures were maintained for 72h. All animal procedures were approved by the UC Davis Institutional Animal Care and Use Committee.

### 2.7 Mouse Liver Tissue Preparation

Male apolipoprotein E-deficient (ApoE^-/-^) mice (B6.129P2-Apoe^tm1Unc^/J; The Jackson Laboratory, Strain #002052) on a C57BL/6J background, a well-established model of diet-induced dyslipidemia and hepatic steatosis, were used in this study. At 8 weeks of age, mice were started on dietary intervention and maintained for 12 weeks. Animals were randomly assigned to receive either a high-fat diet (HFD; 40% kcal from fat; Research Diets, D12109C) or standard laboratory chow, both provided ad libitum. After 12 weeks on diet, mice were euthanized for tissue harvesting. Livers were excised, briefly rinsed in ice-cold PBS to remove residual blood, and immediately processed as fresh tissue for multimodal imaging using stimulated Raman scattering (SRS) and fluorescence lifetime imaging microscopy (FLIM), which provide label-free chemical and metabolic contrast. Samples 1 to 6 corresponded to ApoE^-/-^ mice fed on HFD, whereas samples 7 to 12 corresponded to ApoE^-/-^ mice fed on standard chow. Three animals per group were used for the multimodal imaging analyses reported here (Fig. 3); the imaged subset was selected before analysis and without reference to imaging outcomes. The ROIs of the subsets had enough distance with each other to represent characteristics of wide ROIs. Imaging acquisition and quantitative analyses were performed with investigators blinded to group allocation. All animal procedures were approved by the Institutional Animal Care and Use Committee of the University of California San Diego (protocol S12263) and conducted in accordance with institutional and national guidelines.

**Fig. 3.**
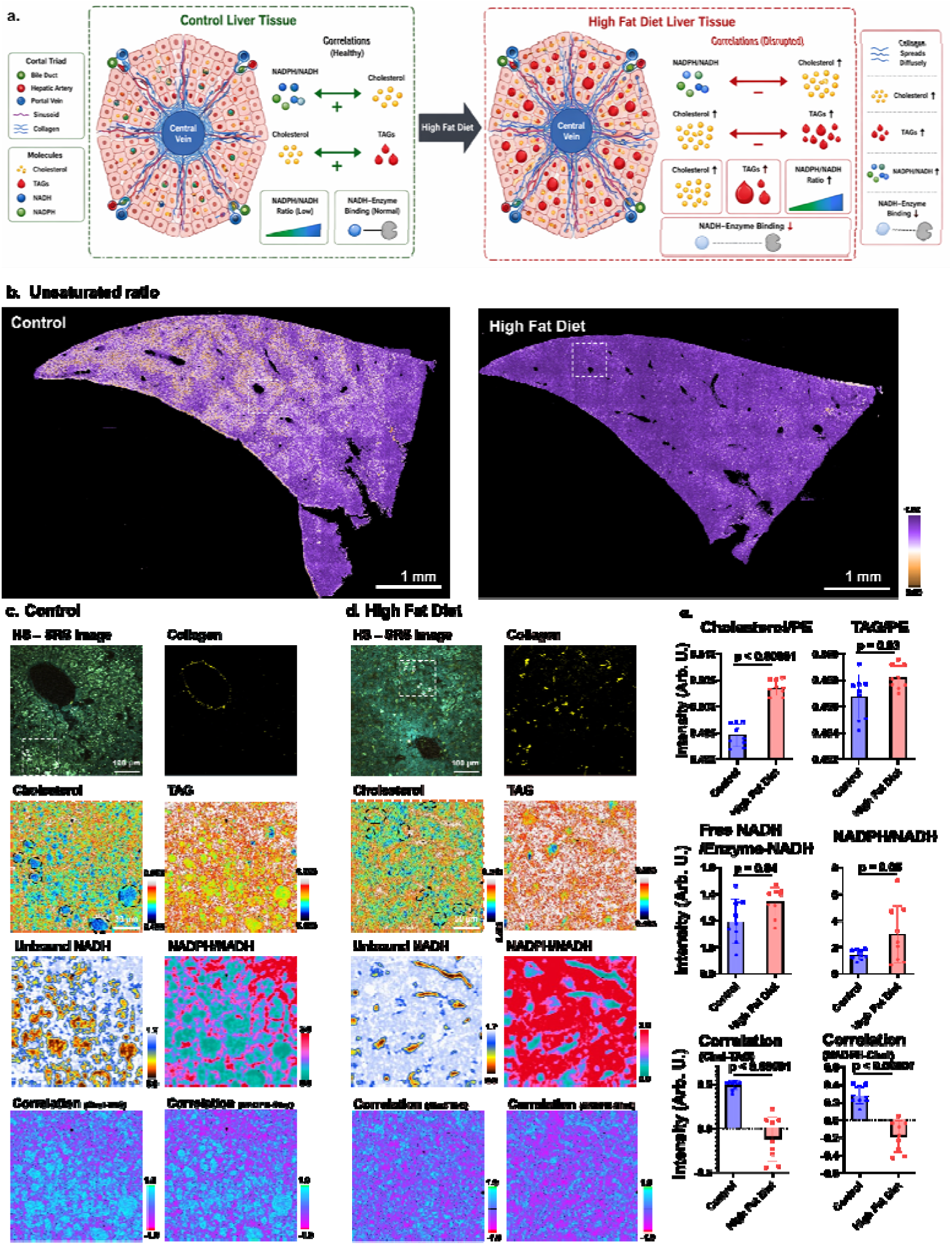
High-fat diet disrupts lipid zonation and alters metabolic and structural organization in mouse liver. **a.** Schematic summarizes the main information about high-fat diet induced liver metabolic changes that were measured with MANIFEST. **b.** Ratiometric SRS images of unsaturated and saturated lipids in control and high-fat-diet liver tissue. In control liver, unsaturated lipid gradients are evident near veins, whereas these gradients are diminished in high-fat-diet samples, indicating disrupted spatial lipid organization. **c and d.** Multimodal images acquired from the boxed regions in b. Hyperspectral SRS images were used to quantify cholesterol and triacylglycerol (TAG) in lipid-rich regions. FLIM images report NADH-associated enzyme-binding activity in the same regions. SHG images show collagen distribution within the tissue. White circles in the HS-SRS images and black circles in the cholesterol images indicate representative regions used for local analysis. **e.** Quantification of multimodal imaging readouts. High-fat diet was associated with increased cholesterol and TAG accumulation together with reduced NADH-associated enzyme-binding activity. Data were acquired from three independent liver samples per group, with three ROIs imaged per sample. ROI-level measurements from the same liver sample were averaged before statistical comparison to yield one value per animal (n = 3 biological replicates per group; 9 ROIs in total). Data are presented as mean ± s.e.m.

### 2.8 Mouse Retina Tissue Slice Preparation

All animal procedures were conducted with the approval of the Institutional Animal Care Committee (IACUC) at the University of California, Irvine, under AUP #23-074. For lipid supplementation studies, male C57BL/6J mice, aged 6 and 26 months, were acquired from Jax Laboratory. The mice were housed in the vivarium at the University of California, Irvine, under a standard 12-hour light (<150 lux)/12-hour dark cycle. They were provided with a standard soy protein-free rodent chow diet (Envigo Teklad 2020X) *ad libitum*.

For D_2_O probing experiments, mice that consumed 20% D_2_O for 7 days were anesthetized using isoflurane and subsequently euthanized. Following euthanasia, mouse eyes were enucleated and immersed in 4% paraformaldehyde (PFA) dissolved in phosphate-buffered saline (PBS, pH 7.4) for 2 hours at 4°C. The cornea, lens, and vitreous were then meticulously removed, and the remaining eyecups were fixed in 4% PFA overnight at 4°C. To facilitate cryosectioning, eyecups underwent cryoprotection by sequential immersion in a sucrose gradient: 10% and 20% sucrose for 1 hour each at room temperature, followed by 30% sucrose overnight at 4°C. The cryoprotected eyecups were then embedded in Tissue-Tek OCT (Sakura, Torrance, CA) and rapidly frozen on a conductive metal block cooled by dry ice. Cryosectioning was performed using a cryostat to obtain 10 µm-thick serial sections of the retina.

### 2.9 Ethical Approval for Human Specimens

The study is compliant with all relevant ethical regulations and was approved by the University of Pennsylvania Institutional Review Board (IRB protocol #848421). Informed consent was obtained from each patient prior to tissue collection by the Human Heart Tissue Bank, and no compensation was provided for participation. Human heart tissue samples were obtained through the Human Heart Tissue Bank from two groups: non-failing, brain-dead organ donors with no clinical history of heart failure (Donor) and patients with end-stage heart failure undergoing transplant (NICM). Heart tissue procurement followed a standardized protocol as previously described[44].

### 2.10 SHG Collagen Analysis

Collagen signals in SHG images were segmented using intensity thresholding, and collagen density was defined as the area fraction of collagen-positive pixels within each ROI. Fiber thickness and orientation were quantified by Otsu thresholding and orientation tensor analysis. The alignment metric reported in the supplementary figures reflects the angular dispersion of local fiber orientations, with lower values indicating a more ordered and parallel fiber architecture. SHG intensities were compared only among samples imaged under identical acquisition settings, and no correction was applied for polarization dependence or imaging depth. Accordingly, “collagen density” refers to area fraction rather than absolute collagen concentration.

### 2.11 Statistics

All quantitative data are presented as mean ± s.e.m., unless otherwise indicated. Statistical analyses were performed using GraphPad Prism 10 (GraphPad Software) and Python. The biological replicate was defined as an independent culture preparation, animal, or human tissue sample, depending on the experiment. Multiple regions of interest (ROIs) acquired from the same biological sample were treated as technical sampling units used to improve spatial representation of that sample rather than as independent biological replicates. Accordingly, unless otherwise stated, measurements derived from multiple ROIs within the same biological sample were first averaged to generate a single value per biological replicate for statistical comparison between groups.

For lipid-droplet (LD)-level analyses, individual droplets were segmented and quantified as described above. Because LDs obtained from the same ROI or biological sample are not statistically independent, droplet-level measurements were used for distributional visualization and descriptive analysis, whereas statistical testing was performed on values summarized at the biological replicate level. The number of biological replicates, ROIs, and segmented objects analyzed for each experiment is reported in the corresponding figure legends.

For comparisons between two groups, two-tailed unpaired Student’s t-tests were used for normally distributed data. For data that did not satisfy normality assumptions, two-tailed Mann–Whitney tests were used. For comparisons involving more than two groups, one-way analysis of variance (ANOVA) followed by Tukey’s multiple-comparisons test was used for parametric data, whereas Kruskal–Wallis tests followed by Dunn’s multiple-comparisons test were used for nonparametric data. Exact n values are provided in the figure legends. Differences were considered statistically significant at P < 0.05.

## 3. Results

### 3.1 Concept and analytical workflow of MANIFEST

To enable integrated, label-free analysis of metabolic heterogeneity, we built MANIFEST, a multimodal nonlinear imaging platform that combines SRS, SHG, MPF, and FLIM on a single microscope framework (Fig. 1a, b, and c). In this configuration of Fig. 1b, pump and Stokes beams are delivered to the sample in a confocal arrangement to generate SRS signals, which are detected in transmission, whereas back-propagating photons are spectrally separated and directed to detectors for SHG, MPF, and FLIM measurements. The Stokes beam is used to generate SHG while minimizing two-photon excitation of NADH, whereas the pump beam enables two-photon excitation of both NADH and FAD. Images from the different modalities were acquired sequentially from the same regions of interest. Collected FLIM images were analyzed using a two-step process of phasor analysis and exponential decay curve fitting (Fig. S1). After phasor analysis[23], every pixel was clustered into a small number of groups with similar lifetime decay curves. The decay curves were averaged, and a representative decay curve was generated from the average. By fitting the curves with biexponential model, fluorescence lifetime values of each group were calculated. From the clustering and initial fitting process, initial values for precise decay curve fitting were prepared. After finding initial values for the fitting, pixel-by-pixel curve fitting was done.

Different analytical approaches were used for images of SRS, SHG, and MPF (Fig. 1c). SRS images were taken as a form of hyperspectral images. After taking hyperspectral images in the C-H stretching region (wavenumber 2700 – 3120 cm^-1^) and cell silent region (wavenumber 2000 – 2300 cm^-1^), each representative image channel was extracted and divided into various metabolic activity images, such as relative protein and lipid turnover. For precise HS-SRS image analysis in the C-H stretching region, the water signal was removed by subtracting the Raman spectrum of the PBS buffer (Fig. S2a). Following this subtraction, lipids were quantified by calculating the lipid-to-water ratio, using a method similar to a previously established approach[17] (Fig. S2b). In addition, integrating Penalized Reference Matching algorithm with SRS (PRM-SRS)[18] for hyperspectral image analysis, lipid subtype distribution was investigated. For MPF and SHG images, intensity-based direct analysis methods were used as in previous studies[34, 35, 45].

After the analysis of each individual channel of biomolecules and metabolic activities, their spatial correlations were calculated. By calculating Pearson’s correlation coefficients of small patches of images (Fig. 1d), spatial correlation mapping is available. As demonstrated by simulated dataset (Fig. S3), the simple calculation of Pearson’s coefficients mapping can show the information of the signals of increasing together, decreasing together, or changing in opposite directions. If two signals increase or decrease together at the same position, their spatial correlation will be positive. If the signals change in opposite directions with each other, then their spatial correlation will be negative values. Using the approach, locally correlated metabolic signals, for example, correlation between NADPH/NADH ratio and lipid amount can be detected (Fig. 1e).

### 3.2 Metabolic alterations in brain cells treated with Amyloid-beta

We first applied MANIFEST to amyloid-beta (Aβ)-treated tri-cultured brain cells as an organelle-level model of neurodegeneration-associated metabolic remodeling. This tri-culture system, containing neurons, astrocytes, and microglia, allowed us to test whether MANIFEST could resolve coordinated changes in lipid composition and cofactor-associated metabolism within individual lipid droplets and their surrounding cellular environment (Fig. 2a). Using hyperspectral SRS together with MPF and FLIM (Fig. 2b and c), we quantified lipid subtype distributions, redox-associated readouts, and cofactor-state changes in control and Aβ-treated cultures [43] (Fig. 2c). HS-SRS microscopy provided insights into the chemical composition of lipid structures, specifically, the ratios of unsaturated lipids, triacylglycerol (TAG), cholesterol ester, and lipid amount. MPF and FLIM of NADH and FAD were used to assess key metabolic indicators such as the redox state, the unbound NADH ratio, and the NADPH ratio, offering a comprehensive view of cellular metabolic activity with their spatial correlations (Fig. 2c).

Detailed examination of individual lipid droplets (LDs) revealed significant heterogeneity and distinct alterations induced by Aβ treatment (Fig. 2d). In the Aβ-treated cells, LDs demonstrated a notable shift in their composition, characterized by an increase in TAG and cholesterol ester content. This compositional change was accompanied by a clear shift in the metabolic state. Specifically, these LDs exhibited a significantly higher redox ratio, a lower unbound NADH ratio, and an elevated NADPH ratio (Fig. 2d and e). The elevated signals of total lipids, cholesterol ester, and NADPH ratio showed distinct spatial correlation patterns: the spatial distribution of lipid amount was more negatively correlated with both cholesterol ester and NADPH ratio in Aβ-treated cells. These metabolic shifts collectively indicate an oxidized and stressed cellular environment, highlighting the detrimental impact of Aβ on brain cell metabolism and lipid homeostasis.

### 3.3 High fat diet induced metabolic changes in mouse liver

We next applied MANIFEST to mouse liver tissue to determine whether multimodal imaging could resolve diet-associated metabolic remodeling at the tissue-compartment level. In control liver, ratiometric SRS imaging revealed clear spatial gradients of unsaturated lipids near veins, consistent with organized metabolic zonation. In contrast, these gradients were diminished in liver tissue from mice fed a high-fat diet, suggesting disruption of normal spatial lipid organization (Fig. 3a and b).

Further analysis revealed more detailed diet-induced changes at the molecular level. HS-SRS imaging showed that the high-fat diet led to a greater accumulation of both cholesterol and TAG within lipid-dense areas. FLIM provided insights into metabolic enzyme activity, indicating that the high-fat diet resulted in reduced enzyme-binding activity for NADH. Additionally, SHG imaging revealed that collagen was widely distributed near hepatocytes, but at a lower density in high-fat diet samples (Fig. S4). These combined findings illustrate that a high-fat diet not only increases lipid storage but also induces significant metabolic stress and structural changes associated with liver disease (Fig. 3c-e). The positive spatial correlations between metabolic signals, such as cholesterol–TAG and NADPH ratio– cholesterol, shifted to negative correlations under the high-fat diet.

### 3.4 Metabolic alteration due to non-ischemic cardiomyopathy

To assess the utility of MANIFEST in human disease tissue, we next examined metabolic and compositional heterogeneity in cardiac samples from non-failing donor hearts and hearts with non-ischemic cardiomyopathy (NICM) (Fig. 4a). SRS imaging revealed distinct spatial distributions of lipids in healthy and diseased myocardium, with higher-magnification images showing marked heterogeneity among lipid-rich structures within individual cardiac cells (Fig. 4b and c).

**Fig. 4.**
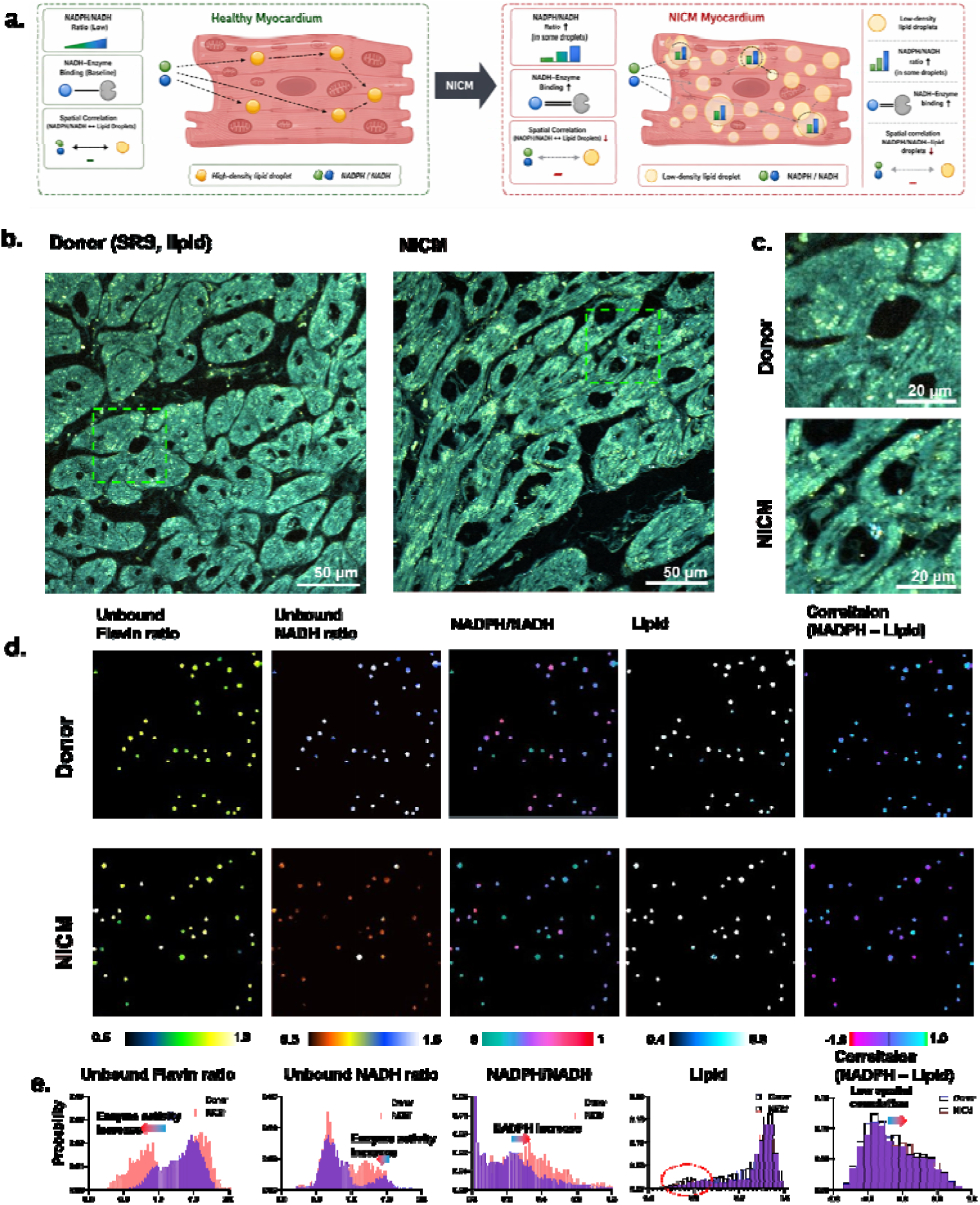
Non-ischemic cardiomyopathy is associated with intracellular lipid remodeling and extracellular matrix reorganization in human heart tissue. **a.** Schematic summarizes the main information about NICM induced metabolic change in lipid droplets that were measured with MANIFEST. **b.** SRS images showing lipid distribution in donor and non-ischemic cardiomyopathy (NICM) cardiac tissue. Images were acquired from three independent human cardiac tissue samples per group, with five ROIs imaged per sample. ROIs from the same tissue sample were treated as technical sampling units. **c.** Enlarged views of the boxed regions in a, highlighting lipid heterogeneity within individual cardiac cells. **d.** High-magnification views of lipid droplets (LDs) from the boxed regions in c. The four channels show unbound flavin ratio, unbound NADH ratio, NADPH-associated signal, and total lipid amount, revealing heterogeneity among individual LDs. **e.** Quantification of LD-associated compositional and metabolic parameters in donor and NICM tissue. Compared with donor tissue, LDs in NICM samples showed reduced unbound flavin and unbound NADH ratios, increased NADPH-associated signal, and reduced lipid signal. Quantification of the averaged SRS lipid channel further identified a subset of LDs with lower lipid content in NICM tissue (red dotted circle in histogram). For group comparisons, measurements from multiple ROIs within the same tissue sample were averaged to generate one value per biological replicate before statistical analysis (n = 3 human tissue samples per group). LD counts are shown for descriptive distributional analysis only and were not treated as independent biological replicates (7,398 LDs in donor tissue; 2,716 LDs in NICM tissue). Data are presented as mean ± s.e.m.

A deeper, quantitative analysis of individual lipid droplets using FLIM uncovered specific metabolic and compositional shifts associated with NICM (Fig. 4d and e). Within the LDs of NICM tissue, there was a notable decrease in the ratios of unbound Flavin and unbound NADH. This suggests that a larger fraction of these essential metabolic coenzymes is enzyme-bound, indicating a potential alteration in cellular energy metabolism. Concurrently, an increase in NADPH level was observed within these same LDs, which may point to changes in biosynthetic and antioxidant pathways. In addition, the spatial correlation between NADPH level and lipid amount became more heterogeneous in NICM. Compositionally, the analysis also revealed a decrease in lipid amount within the LDs of NICM tissue. In addition, SHG imaging revealed marked changes in collagen distribution, which was more widely distributed and exhibited lower angular dispersion (i.e., more ordered fiber alignment) in NICM tissues (Fig. S5).

### 3.5 Metabolic activity changes in aging mouse retina

Finally, we applied MANIFEST to the aging mouse retina to investigate how metabolic heterogeneity is reorganized across anatomically distinct retinal compartments with age. The retina is a highly structured tissue in which different layers have specialized functions, lipid composition, and metabolic demand. We therefore compared retinal tissues from young (6-month) and aged (26-month) mice using FLIM/MPF, SRS, SHG, and DO-SRS to map age-associated changes in cofactor state, lipid composition, collagen, and lipid turnover (Fig. S6). Pseudo-H&E images were used to identify key compartments, including the outer nuclear layer (ONL), retinal pigment epithelium (RPE), and rod outer segments (Fig. S7). Ratiometric analyses generated from MANIFEST revealed compartment-specific differences in the balance of free and enzyme-bound NADH, the NADPH/NADH ratio, the sphingosine-to-phosphatidylethanolamine (PE) ratio, and relative lipid turnover (Fig. 5a). These images demonstrate that different layers of the retina, such as the rod cells and the retinal pigment epithelium (RPE), exhibit unique metabolic signatures that change significantly with age.

**Fig. 5.**
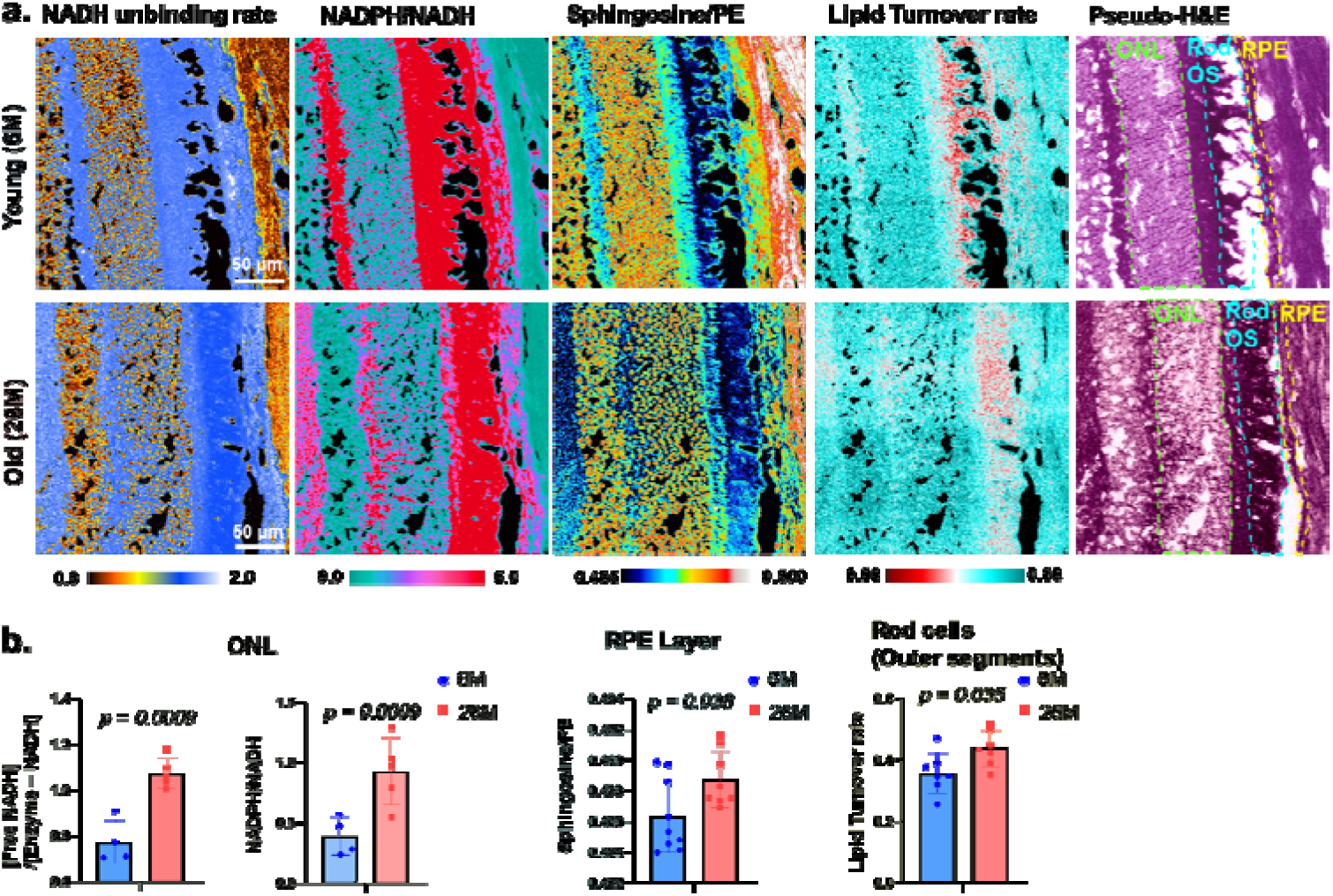
Aging remodels cofactor state, lipid composition, and lipid turnover across retinal compartments. **a.** Ratiometric images generated by MANIFEST from retinal tissue of young (6-month) and aged (26-month) mice. Pseudo-H&E images were used to define the outer nuclear layer (ONL; green dashed region), retinal pigment epithelium (RPE; yellow dashed region), and rod outer segments (cyan dashed region). The corresponding maps show compartment-resolved differences in free/enzyme-bound NADH, NADPH/NADH, sphingosine/PE, and relative lipid turnover. **b.** Quantification of compartment-specific metabolic and compositional differences between young and aged retina. FLIM analysis showed altered free NADH-associated signal in the ONL of aged mice. DO-SRS revealed differences in lipid turnover in rod outer segments with age. PRM-SRS analysis showed age-associated differences in sphingosine relative to phosphatidylethanolamine (PE) in the RPE. For FLIM, DO-SRS, and PRM-SRS analyses, one ROI was analyzed per sample, so n corresponds directly to the number of biological replicates. Sample numbers were: FLIM, n = 5 per age group; DO-SRS, n = 7 per age group; PRM-SRS, n = 9 young and n = 8 aged samples. Data are presented as mean ± s.e.m.

Statistical analysis of the imaging data confirms clear, age-driven differences in the retina’s metabolic and chemical makeup (Fig. 5). FLIM imaging revealed a shift in the proportion of free NADH and NADPH in the outer nuclear layer of old mice, suggesting shifts in cellular energy metabolism. SRS imaging integrated with heavy water (D_2_O) treatment (DO-SRS) revealed changes in relative lipid turnover (D-label incorporation) in the rod cells of old mice, indicating altered lipid dynamics in photoreceptors. Furthermore, PRM-SRS results point to a change in sphingosine within the RPE layer. These findings collectively illustrate a complex landscape of metabolic and structural reorganization in the aging retina.

## 4. Discussion

In this study, we developed MANIFEST, a label-free multimodal imaging platform that integrates stimulated Raman scattering (SRS), second harmonic generation (SHG), multiphoton fluorescence (MPF), and fluorescence lifetime imaging microscopy (FLIM) to map metabolic heterogeneity with their spatial correlations across biological scales. By combining SRS-based chemical imaging of lipids with autofluorescence- and lifetime-based analysis of endogenous cofactors, MANIFEST links macromolecular composition, redox-associated metabolism, and structural organization within the same specimen. Across four distinct biological contexts—amyloid-beta-treated brain tri-cultures, high-fat-diet liver, human non-ischemic cardiomyopathy tissue, and aging retina—this framework resolved coordinated but spatially restricted changes in lipid remodeling, cofactor state, collagen architecture, and tissue-compartment organization. These results highlight the value of multimodal nonlinear imaging not only for detecting biochemical differences, but also for placing those differences into their spatial and structural context, from organelle-associated lipid droplets to intact tissue compartments.

A central technical contribution of MANIFEST is the integration of the FLIM modality, together with a spatial correlation analysis pipeline, into prior multimodal nonlinear optical imaging strategies. Previous multimodal platforms incorporating SRS, SHG, and MPF have demonstrated strong capabilities for visualizing lipid distributions, extracellular matrix structure, and endogenous fluorescence signals across cells and tissues [34, 35, 45]. Here, we extend that framework by incorporating spatial correlation mapping method and phasor-guided, biexponential FLIM analysis to distinguish cofactor-associated fluorescence states of NADH and FAD and to relate these signals directly to surrounding lipid-rich structures and tissue features. This addition of FLIM analysis is important because metabolic cofactors report biochemical processes that are not directly accessible through macromolecular imaging alone. By integrating SRS-derived chemical contrast with FLIM-based cofactor-state information, MANIFEST expands label-free metabolic imaging from compositional analysis toward a more functionally informative view of cellular state. In addition, because the biochemical reactions can support or compete with each other, the spatial correlation mapping method can boost the advantage of the multimodal imaging approach.

The multimodal structure of MANIFEST is particularly valuable because similar molecular readouts can arise from distinct underlying biological mechanisms. This point is illustrated across the four model systems examined here. In amyloid-beta-treated brain cells, changes in lipid droplet composition were accompanied by shifts in redox ratio, unbound NADH, and NADPH-associated signals, consistent with stress-associated metabolic remodeling in a neuroinflammatory context. In the high-fat-diet liver, altered lipid accumulation occurred together with disrupted spatial zonation and changes in collagen organization, indicating that metabolic remodeling extended beyond intracellular lipid storage to tissue-level reorganization. In non-ischemic cardiomyopathy, elevated NADPH-associated signal was observed despite reduced lipid signal within lipid droplets, underscoring that an increase in NADPH-related fluorescence does not necessarily imply increased lipid anabolism. Instead, such a shift may reflect other processes, including oxidative stress adaptation or altered enzyme activity. These examples illustrate why multimodal readouts are needed: the same apparent metabolic signature may have different biological meanings depending on the surrounding chemical and structural context. Conversely, no subset of the modalities would support these interpretations on its own: SRS provides the chemical identity and subcellular localization that anchor the analysis, FLIM disambiguates cofactor pools that are indistinguishable by intensity alone, MPF supplies the redox-ratio readout, and SHG places both in the context of extracellular matrix remodeling. Removing any one channel would leave the corresponding axis of interpretation unconstrained.

The amyloid-beta tri-culture dataset provides proof of concept for organelle-level metabolic phenotyping with MANIFEST. In this system, hyperspectral SRS and FLIM revealed that individual lipid droplets are compositionally and metabolically heterogeneous and that this heterogeneity is reshaped by amyloid-beta exposure. The observed increase in TAG and cholesterol ester signals, together with altered cofactor-associated readouts, is consistent with the growing view that lipid-droplet remodeling is closely linked to neuroinflammation and neurodegeneration [46–48]. Importantly, MANIFEST resolves these changes at the level of individual droplets rather than only at the whole-cell level, demonstrating its utility for studying organelle-associated metabolic states in complex cellular models. More broadly, this result highlights how multimodal label-free imaging can connect lipid remodeling with changes in endogenous cofactor behavior within the same subcellular structures[47, 48].

The liver data demonstrates that MANIFEST can resolve metabolic remodeling at the level of tissue organization. In control liver, ratiometric SRS imaging revealed spatial gradients of unsaturated lipids near veins, reflecting the well-known zonated structure of hepatic metabolism. Under high-fat-diet conditions, these gradients were diminished, indicating disruption of normal metabolic patterning. This result is especially important because it shows that metabolic dysfunction in liver disease is not only a matter of increased lipid storage, but also a loss of spatial organization. The accompanying changes in collagen signal detected by SHG further suggest that metabolic and structural remodeling occur together in diet-induced liver pathology. In this context, MANIFEST provides a way to bridge intracellular lipid chemistry with tissue-scale organization, which is difficult to achieve using either bulk biochemical assays or single-modality imaging alone.

The analysis of human NICM extends this framework to clinically relevant tissue. Cardiomyopathy involves remodeling of energy metabolism, lipid handling, and extracellular matrix composition, all of which are spatially heterogeneous within the myocardium. MANIFEST resolved differences in lipid-droplet-associated cofactor states together with pronounced collagen reorganization in NICM tissue, showing that diseased myocardium differs from donor tissue at both intracellular and extracellular levels. The ability to analyze endogenous metabolic and compositional features directly in human tissue without labeling is a notable strength of the approach. Given the small number of donors examined here, these observations are exploratory and will require validation in larger cohorts; nevertheless, they suggest that MANIFEST may be useful for identifying disease-associated metabolic phenotypes in human biopsy or surgical tissue samples, particularly in contexts where tissue heterogeneity is central to pathology.

Application of spatial correlation mapping across three different sample types highlights a unique strength of MANIFEST. Even when multiple metabolic activities are simultaneously elevated, the correlations between them can vary substantially. A representative example is the relationship between NADPH and lipids. In this study, correlation mapping was primarily applied to the NADPH ratio in relation to total lipid content or specific lipid subtypes. In tri-cultured cell samples, the spatial correlations between the NADPH ratio and lipid content, as well as between cholesterol ester and total lipid, were negative. Because NADPH is consumed during lipid synthesis[49], the local NADPH ratio may show a negative correlation with lipid abundance, even when the overall NADPH pool is elevated to support active lipogenesis. In addition, since the total lipid signal mainly reflects fatty acid-derived signals, the negative correlation between total lipid content and cholesterol ester likely indicates competition for the limited space within lipid droplets[50]. A similar spatial correlation pattern was also observed in the liver images. Furthermore, the correlation changes identified in NICM heart tissue demonstrate that this approach can be used to reveal shifts in metabolic roles under altered chemical conditions and disease states.

The aging retina provides a further example of how multimodal imaging can reveal compartment-specific metabolic remodeling in a highly structured tissue. The retina is composed of layers with distinct metabolic roles, and age-associated dysfunction is likely to emerge unevenly across these compartments. MANIFEST captured such differences by resolving altered cofactor-associated states in the outer nuclear layer, altered lipid turnover in rod outer segments, and changes in lipid subtype composition in the retinal pigment epithelium. These data underscore that retinal aging is spatially organized rather than uniform and illustrate the advantage of integrating FLIM, PRM-SRS, and DO-SRS within a single framework (Fig. S7). More broadly, the retina results highlight one of the central strengths of MANIFEST: its ability to connect biochemical composition, cofactor-state information, and tissue architecture in tissues where functional specialization is tightly coupled to spatial organization.

Another important strength of MANIFEST is its ability to operate across scales. At the subcellular level, it resolves lipid-droplet heterogeneity and links droplet composition to cofactor-associated metabolic signals. At the tissue level, it reveals compartment-specific metabolic differences and matrix remodeling. This multiscale capability is particularly relevant for studies of aging and disease, where pathology often emerges through coupled changes in intracellular metabolism, extracellular structure, and tissue organization. Because the platform is label-free and relies on endogenous contrasts, it is well suited for comparative studies across diverse biological systems and can be integrated with existing models of disease progression and tissue dysfunction.

At the same time, several limitations should be noted. First, the different imaging modalities are acquired sequentially rather than simultaneously, which may limit throughput and introduce practical constraints for highly dynamic live-cell applications. Sequential acquisition also carries a risk of sample drift between modalities; in the present study all modalities were acquired from the same fields of view in immediate succession to minimize drift, but simultaneous or interleaved acquisition schemes will be needed to extend MANIFEST to fast dynamics in living systems. Second, although FLIM-derived cofactor-state measurements provide valuable functional information, the biological interpretation of these signals remains context dependent and should not be reduced to a single pathway-level readout. In particular, oxidative stress can manifest as shifts in either NAD(P)H- or FAD-associated signals, so the sign of a change in the optical redox ratio between control and pathological samples is not by itself diagnostic of a specific mechanism; perturbation experiments with defined metabolic challenges will be required to validate these cofactor-state interpretations. As shown here, changes in NADPH-associated signals can reflect different underlying processes across tissues. Third, while MANIFEST provides strong correlative information linking composition, cofactor state, and structure, it does not by itself establish mechanism. Follow-up perturbation experiments, isotope tracing, or orthogonal biochemical assays will be important for testing the causal basis of the metabolic states observed here. Finally, because the current implementation emphasizes lipids and endogenous cofactors, future extensions could expand the molecular scope further through improved hyperspectral analysis, additional vibrational contrasts, or integration with complementary spatial omics approaches. The present analyses also relied on univariate comparisons of individual readouts complemented by pairwise spatial correlation mapping; integrating the co-registered multimodal channels into multivariate or machine learning-based phenotyping, together with cell segmentation that assigns metabolic signatures to specific cell types and functional tissue units, is an important next step that these datasets are well suited to enable.

In summary, these proof-of-concept studies establish MANIFEST as an extensible framework for label-free, high-resolution metabolic phenotyping in complex biological systems. By integrating SRS, SHG, MPF, and FLIM within a unified analytical platform, it reveals how lipid composition, cofactor-associated metabolism, and tissue structure are spatially coordinated across organelles, cells, and tissue compartments. Across disease and aging models, this approach resolved metabolic heterogeneity that would be difficult to detect using any single modality alone. These capabilities position MANIFEST as a versatile tool for studying metabolism-driven tissue pathology and for advancing mechanistic investigation of aging and disease in situ.

## Acknowledgements

We thank Wei Min, Christian Metallo, Uri Alon, Peter Adams, Rob Knight, and Shi Lab members for meaningful discussions and feedback.

## 7. Author Contributions

L. S. conceived the idea and designed the project; H. J. imaged tri-cultured cells, cardiac tissues, and retina tissues; S. W. imaged liver tissues and retina tissues; H.J. analyzed the images and generated figures with the input from L.S.; H. K. and E. S. prepared tri-cultured cell samples; F. Y. D.S. prepared mice retina samples; T.W.W, J.K, W.S, and J.S prepared mice liver samples; A.A and A.J prepared human cardiac samples; H. J. and L.S. wrote and revised the manuscript with the input from all other authors.

## 8. Data availability statement

Scripts for data analysis were written mostly in Python, available at https://github.com/lingyanshi2020/MANIFEST. Experimental data will be provided as requested.

## 9. Supplementary Document

See Supplement 1 for supporting figures.

## References

1. J. Villazon, Z. Li, A. Fan, and L. Shi, “Multimodal Optical Imaging Platform for Studying Cellular Metabolism,” JoVE, e67906 (2025).

2. J. W. Lichtman, and J.-A. Conchello, “Fluorescence microscopy,” Nature Methods 2, 910–919 (2005).

3. N. de Souza, S. Zhao, and B. Bodenmiller, “Multiplex protein imaging in tumour biology,” Nature Reviews Cancer 24, 171–191 (2024).

4. S. W. Hell, and J. Wichmann, “Breaking the diffraction resolution limit by stimulated emission: stimulated-emission-depletion fluorescence microscopy,” Opt. Lett. 19, 780–782 (1994).

5. E. Betzig, G. H. Patterson, R. Sougrat, O. W. Lindwasser, S. Olenych, J. S. Bonifacino, M. W. Davidson, J. Lippincott-Schwartz, and H. F. Hess, “Imaging intracellular fluorescent proteins at nanometer resolution,” science 313, 1642–1645 (2006).

6. M. J. Rust, M. Bates, and X. Zhuang, “Sub-diffraction-limit imaging by stochastic optical reconstruction microscopy (STORM),” Nature methods 3, 793–796 (2006).

7. M. G. Gustafsson, “Surpassing the lateral resolution limit by a factor of two using structured illumination microscopy,” Journal of microscopy 198, 82–87 (2000).

8. M. Weber, M. Leutenegger, S. Stoldt, S. Jakobs, T. S. Mihaila, A. N. Butkevich, and S. W. Hell, “MINSTED fluorescence localization and nanoscopy,” Nature Photonics 15, 361–366 (2021).

9. T. Dertinger, R. Colyer, R. Vogel, M. Heilemann, M. Sauer, J. Enderlein, and S. Weiss, “Superresolution optical fluctuation imaging (SOFI),” Nano-Biotechnology for Biomedical and Diagnostic Research, 17–21 (2012).

10. A. Ghosh, A. Sharma, A. I. Chizhik, S. Isbaner, D. Ruhlandt, R. Tsukanov, I. Gregor, N. Karedla, and J. Enderlein, “Graphene-based metal-induced energy transfer for sub-nanometre optical localization,” Nature Photonics 13, 860–865 (2019).

11. S. Hell, and E. H. K. Stelzer, “Properties of a 4Pi confocal fluorescence microscope,” J. Opt. Soc. Am. A 9, 2159–2166 (1992).

12. W. Min, C. W. Freudiger, S. Lu, and X. S. Xie, “Coherent Nonlinear Optical Imaging: Beyond Fluorescence Microscopy,” Annual Review of Physical Chemistry 62, 507–530 (2011).

13. V. Parodi, E. Jacchetti, R. Osellame, G. Cerullo, D. Polli, and M. T. Raimondi, “Nonlinear Optical Microscopy: From Fundamentals to Applications in Live Bioimaging,” Frontiers in Bioengineering and Biotechnology **Volume** 8 - 2020 (2020).

14. R. Li, X. Wang, Y. Zhou, H. Zong, M. Chen, and M. Sun, “Advances in nonlinear optical microscopy for biophotonics,” Journal of Nanophotonics 12, 033007 (2018).

15. C. W. Freudiger, W. Min, B. G. Saar, S. Lu, G. R. Holtom, C. He, J. C. Tsai, J. X. Kang, and X. S. Xie, “Label-free biomedical imaging with high sensitivity by stimulated Raman scattering microscopy,” Science 322, 1857–1861 (2008).

16. D. Fu, J. Zhou, W. S. Zhu, P. W. Manley, Y. K. Wang, T. Hood, A. Wylie, and X. S. Xie, “Imaging the intracellular distribution of tyrosine kinase inhibitors in living cells with quantitative hyperspectral stimulated Raman scattering,” Nature chemistry 6, 614–622 (2014).

17. S. Oh, C. Lee, W. Yang, A. Li, A. Mukherjee, M. Basan, C. Ran, W. Yin, C. J. Tabin, D. Fu, X. S. Xie, and M. W. Kirschner, “Protein and lipid mass concentration measurement in tissues by stimulated Raman scattering microscopy,” Proceedings of the National Academy of Sciences 119, e2117938119 (2022).

18. W. Zhang, Y. Li, A. A. Fung, Z. Li, H. Jang, H. Zha, X. Chen, F. Gao, J. Y. Wu, and H. Sheng, “Multi- molecular hyperspectral PRM-SRS microscopy,” Nature Communications 15, 1599 (2024).

19. S. Roth, and I. Freund, “Second harmonic generation in collagen,” The Journal of chemical physics 70, 1637–1643 (1979).

20. P. J. Campagnola, and C. Y. Dong, “Second harmonic generation microscopy: principles and applications to disease diagnosis,” Laser & Photonics Reviews 5, 13–26 (2011).

21. S. Huang, A. A. Heikal, and W. W. Webb, “Two-Photon Fluorescence Spectroscopy and Microscopy of NAD(P)H and Flavoprotein,” Biophysical Journal 82, 2811–2825 (2002).

22. O. I. Kolenc, and K. P. Quinn, “Evaluating Cell Metabolism Through Autofluorescence Imaging of NAD(P)H and FAD,” Antioxidants & Redox Signaling 30, 875–889 (2019).

23. B. Torrado, B. Pannunzio, L. Malacrida, and M. A. Digman, “Fluorescence lifetime imaging microscopy,” Nature Reviews Methods Primers 4, 80 (2024).

24. H. Jang, Z. Li, E. Ackerstaff, J. A. Koutcher, and L. Shi, “Super resolution multimodal microscopy for studying spatially correlated metabolism in aging and diseases,” in Ultrafast Nonlinear Imaging and Spectroscopy XI(SPIE2023), p. 1268102.

25. H. Jang, Y. Li, S. Wu, and L. Shi, “Super-Resolution Stimulated Raman Scattering Microscopy with Graphical User Interface–Supported A-PoD,” Current Protocols 4, e970 (2024).

26. H. Jang, S. Wu, Y. Li, Z. Li, and L. Shi, “Metabolic nanoscopy enhanced by experimental and computational approaches,” npj Imaging 2, 55 (2024).

27. M. C. Skala, K. M. Riching, A. Gendron-Fitzpatrick, J. Eickhoff, K. W. Eliceiri, J. G. White, and N. Ramanujam, “In vivo multiphoton microscopy of NADH and FAD redox states, fluorescence lifetimes, and cellular morphology in precancerous epithelia,” Proceedings of the National Academy of Sciences 104, 19494–19499 (2007).

28. A. A. Heikal, “Intracellular Coenzymes as Natural Biomarkers for Metabolic Activities and Mitochondrial Anomalies,” Biomarkers in Medicine 4, 241–263 (2010).

29. I. Georgakoudi, and K. P. Quinn, “Optical Imaging Using Endogenous Contrast to Assess Metabolic State,” Annual Review of Biomedical Engineering 14, 351–367 (2012).

30. J. Vergen, C. Hecht, L. V. Zholudeva, M. M. Marquardt, R. Hallworth, and M. G. Nichols, “Metabolic Imaging Using Two-Photon Excited NADH Intensity and Fluorescence Lifetime Imaging,” Microscopy and Microanalysis 18, 761–770 (2012).

31. N. Vogler, S. Heuke, T. W. Bocklitz, M. Schmitt, and J. Popp, “Multimodal imaging spectroscopy of tissue,” Annual Review of Analytical Chemistry 8, 359–387 (2015).

32. I. W. Schie, C. Stiebing, and J. Popp, “Looking for a perfect match: multimodal combinations of Raman spectroscopy for biomedical applications,” Journal of biomedical optics 26, 080601–080601 (2021).

33. Y. Lingxiao, P. Jaena, J. C. Eric, E. S. Janet, M. Marina, P. Heidi, R. S. Darold, and A. B. Stephen, “Label-free multimodal nonlinear optical imaging of needle biopsy cores for intraoperative cancer diagnosis,” Journal of Biomedical Optics 27, 056504 (2022).

34. Z. Li, C. Nguyen, H. Jang, D. Hoang, S. Min, E. Ackerstaff, J. A. Koutcher, and L. Shi, “Multimodal imaging of metabolic activities for distinguishing subtypes of breast cancer,” Biomed. Opt. Express 14, 5764–5780 (2023).

35. A. A. Fung, Z. Li, C. Boote, P. Markov, J. P. Gaut, S. Jain, and L. Shi, “Label-free multimodal optical biopsy reveals biomolecular and morphological features of diabetic kidney tissue in 2D and 3D,” Nature Communications 16, 4509 (2025).

36. H. Jang, Y. Li, A. A. Fung, P. Bagheri, K. Hoang, D. Skowronska-Krawczyk, X. Chen, J. Y. Wu, B. Bintu, and L. Shi, “Super-resolution SRS microscopy with A-PoD,” Nature Methods 20, 448–458 (2023).

37. Y. Cheng, “Mean shift, mode seeking, and clustering,” IEEE Transactions on Pattern Analysis and Machine Intelligence 17, 790–799 (1995).

38. D. Xiao, N. Sapermsap, Y. Chen, and D. D. U. Li, “Deep learning enhanced fast fluorescence lifetime imaging with a few photons,” Optica 10, 944–951 (2023).

39. T. S. Blacker, Z. F. Mann, J. E. Gale, M. Ziegler, A. J. Bain, G. Szabadkai, and M. R. Duchen, “Separating NADH and NADPH fluorescence in live cells and tissues using FLIM,” Nature Communications 5, 3936 (2014).

40. T. Blu, and F. Luisier, “The SURE-LET approach to image denoising,” IEEE Transactions on Image Processing 16, 2778–2786 (2007).

41. F. Luisier, T. Blu, and M. Unser, “Image Denoising in Mixed Poisson–Gaussian Noise,” IEEE Transactions on Image Processing 20, 696–708 (2011).

42. S. Bolte, and F. P. CordeliÈRes, “A guided tour into subcellular colocalization analysis in light microscopy,” Journal of Microscopy 224, 213–232 (2006).

43. H. Kim, B. Le, N. Goshi, K. Zhu, A. C. Grodzki, P. J. Lein, M. Zhao, and E. Seker, “Primary cortical cell tri-culture to study effects of amyloid-β on microglia function and neuroinflammatory response,” Journal of Alzheimer’s Disease 102, 730–741 (2024).

44. K. C. Bedi, N. W. Snyder, J. Brandimarto, M. Aziz, C. Mesaros, A. J. Worth, L. L. Wang, A. Javaheri, I. A. Blair, K. B. Margulies, and J. E. Rame, “Evidence for Intramyocardial Disruption of Lipid Metabolism and Increased Myocardial Ketone Utilization in Advanced Human Heart Failure,” Circulation 133, 706–716 (2016).

45. B. L. Gorman, Z. Li, G. Deutsch, H. L. Huyck, N. Beishembieva, H. Bhotika, H. Olson, J. Villazon, P. Yu, G. S. Pryhuber, G. Clair, L. Shi, and C. R. Anderton, “Multiscale metabolic mapping of lung tissue via coregistered mass spectrometry and nonlinear optical imaging,” Science Advances 12, eaec3544 (2026).

46. Y. Li, D. Munoz-Mayorga, Y. Nie, N. Kang, Y. Tao, J. Lagerwall, C. Pernaci, G. Curtin, N. G. Coufal, J. Mertens, L. Shi, and X. Chen, “Microglial lipid droplet accumulation in tauopathy brain is regulated by neuronal AMPK,” Cell Metabolism 36, 1351–1370.e1358 (2024).

47. P. Prakash, P. Manchanda, E. Paouri, K. Bisht, K. Sharma, J. Rajpoot, V. Wendt, A. Hossain, P. R. Wijewardhane, C. E. Randolph, Y. Chen, S. Stanko, N. Gasmi, A. Gjojdeshi, S. Card, J. Fine, K. P. Jethava, M. G. Clark, B. Dong, S. Ma, A. Crockett, E. A. Thayer, M. Nicolas, R. Davis, D. Hardikar, D. Allende, R. A. Prayson, C. Zhang, D. Davalos, and G. Chopra, “Amyloid-β induces lipid droplet-mediated microglial dysfunction via the enzyme DGAT2 in Alzheimer’s disease,” Immunity 58, 1536–1552.e1538 (2025).

48. L. Sánchez de Muniain, P. Escalada, M. J. Ramírez, and M. Solas, “Astrocytes as Metabolic Sensors Orchestrating Energy-Driven Brain Vulnerability in Alzheimer’s Disease,” J Neurochem 169, e70252 (2025).

49. J. Fan, J. Ye, J. J. Kamphorst, T. Shlomi, C. B. Thompson, and J. D. Rabinowitz, “Quantitative flux analysis reveals folate-dependent NADPH production,” Nature 510, 298–302 (2014).

50. S. Yue, J. Li, S.-Y. Lee, Hyeon J. Lee, T. Shao, B. Song, L. Cheng, Timothy A. Masterson, X. Liu, Timothy L. Ratliff, and J.-X. Cheng, “Cholesteryl Ester Accumulation Induced by PTEN Loss and PI3K/AKT Activation Underlies Human Prostate Cancer Aggressiveness,” Cell Metabolism 19, 393–406 (2014).

