## Supplemental Figures for "Label free multimodal optical imaging of metabolic heterogeneity and correlation in aging by integrating SRS, MPF, FLIM, and SHG"

: supplemental document

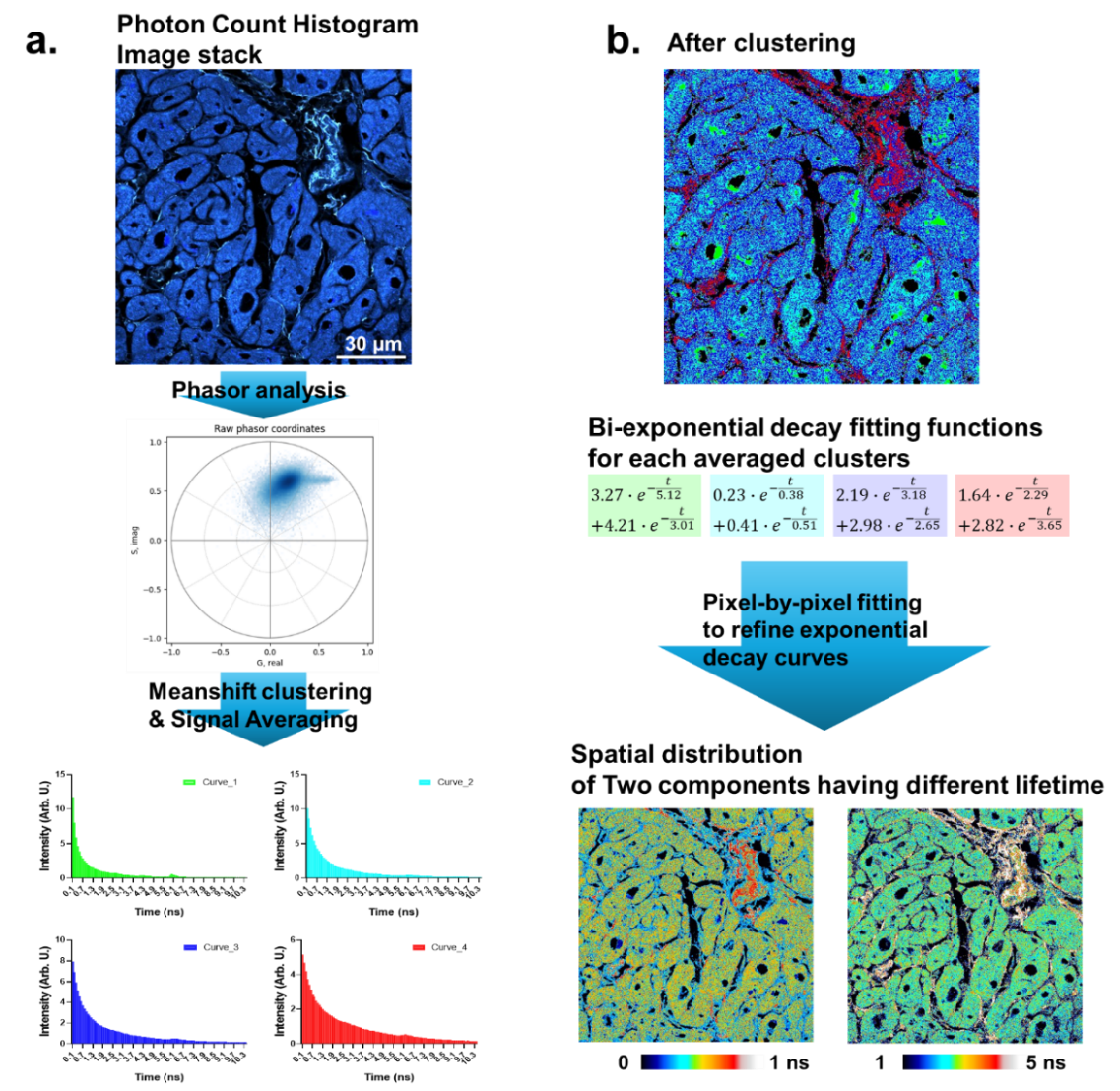

**Fig S1. Schematic of two-step FLIM data analysis a.** In the first step, pixels in a photon counting histogram (PCH) image stack are clustered using the Meanshift algorithm. Within each cluster, the corresponding PCHs are averaged to improve the signal-to-noise ratio prior to further analysis. **b.** In the subsequent step, the averaged PCHs are fitted with a bi-exponential decay function model to extract initial estimates of the fluorescence lifetime components. The fitting parameters are then refined through an additional pixel-by-pixel fitting procedure, in which the PCH of each individual pixel is fitted with the bi-exponential decay function using the cluster-averaged fitting parameters as initial values. This approach ensures robust convergence and improves the accuracy of the fitting at the single-pixel level. From the resulting pixel-wise analysis, the calculated lifetime values of the two components are reconstructed into two spatially resolved lifetime maps, each representing the lifetime of one of the two decay components.

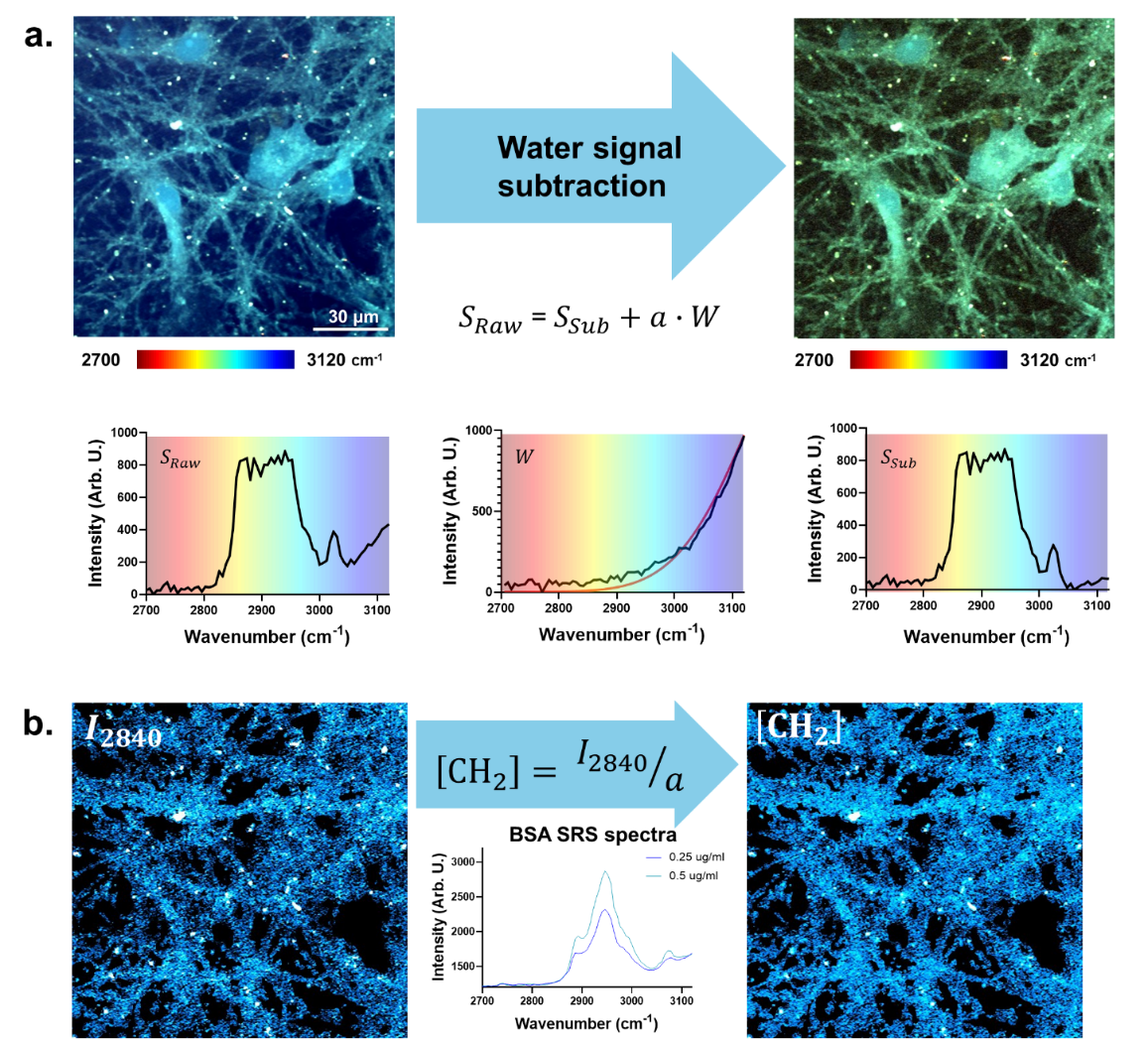

**Fig S2. Quantitative analysis of lipid signal. a.** From a hyperspectral image stack (S_Raw_), the Raman signal of water can be isolated owing to the broad spectral profile of the water Raman band. The Raman spectrum of PBS buffer (W) was acquired separately and subsequently subtracted from S_Raw_ after rescaling by a scaling parameter (*a*), which is proportional to the local water content. The resulting subtracted spectra (S_Sub_) yield a hyperspectral image free from water vibrational contributions. **b.** The -CH_2_ stretching vibration at 2840 cm^-1^ was used to quantify the lipid content in the sample. By normalizing the lipid signal with respect to the water scaling parameter (*a*), absolute amount information was obtained. The validity of this quantification approach was confirmed using BSA spectra acquired at two different concentrations, demonstrating a linear relationship consistent with the expected concentration dependence.

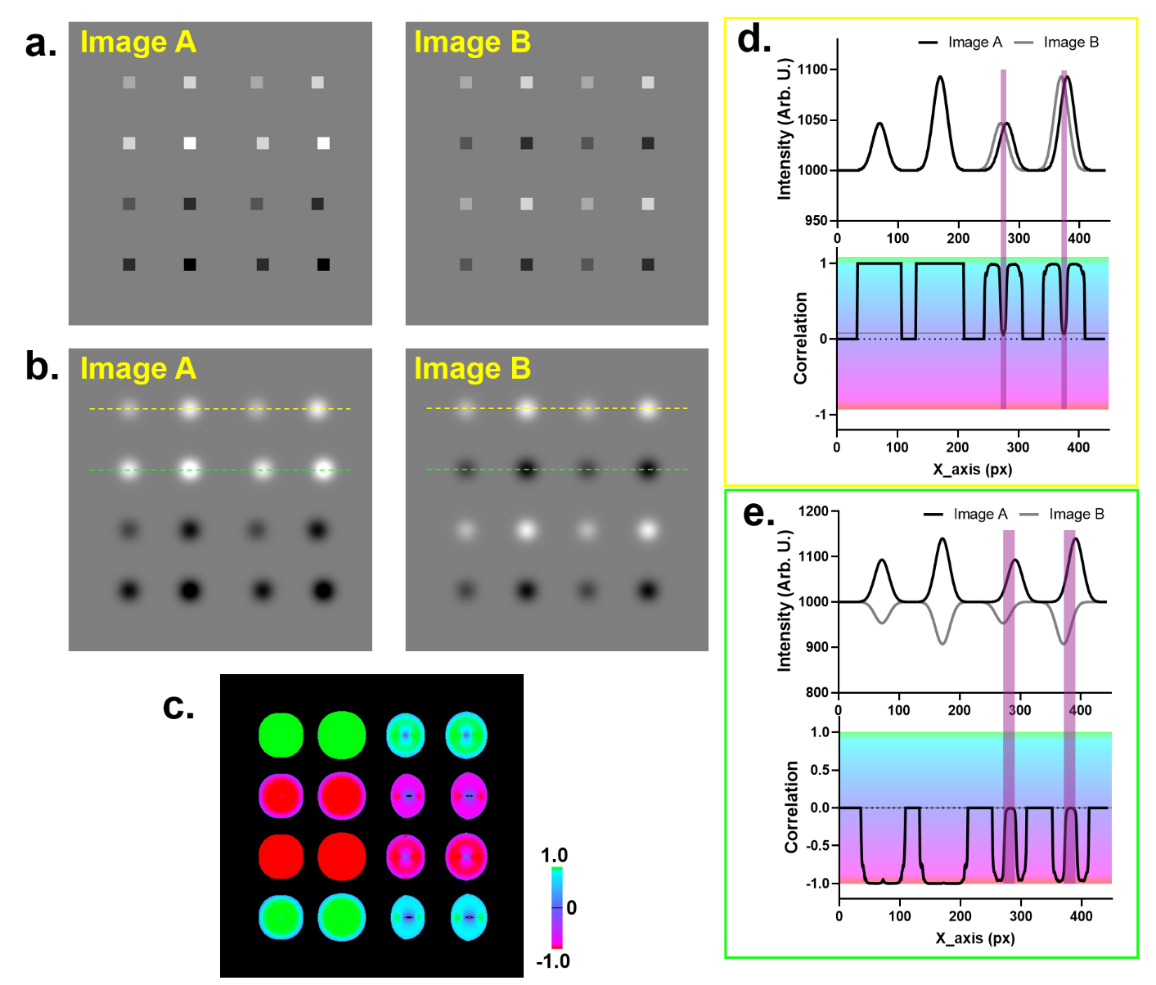

**Fig. S3.** **Spatial correlation mapping simulation.** **a.** Sixteen spots with different intensity values were arranged in a square image. In image B, the second and third rows were assigned intensity values opposite to those of the corresponding rows in image A, and the third and fourth columns were shifted to the right. In these columns, the spots in the second and fourth rows were displaced by twice the distance of those in the first and third rows. **b.** Images A and B were blurred using a Gaussian filter with a sigma of 10 pixels. **c.** The spatial correlation map reveals positively and negatively correlated intensity changes. **d and e.** When intensity changes occur within the same ROIs, strong spatial correlation values are observed. In the purple region of the intensity plots, however, the displacement of the spots causes a mismatch in the intensity gradients.

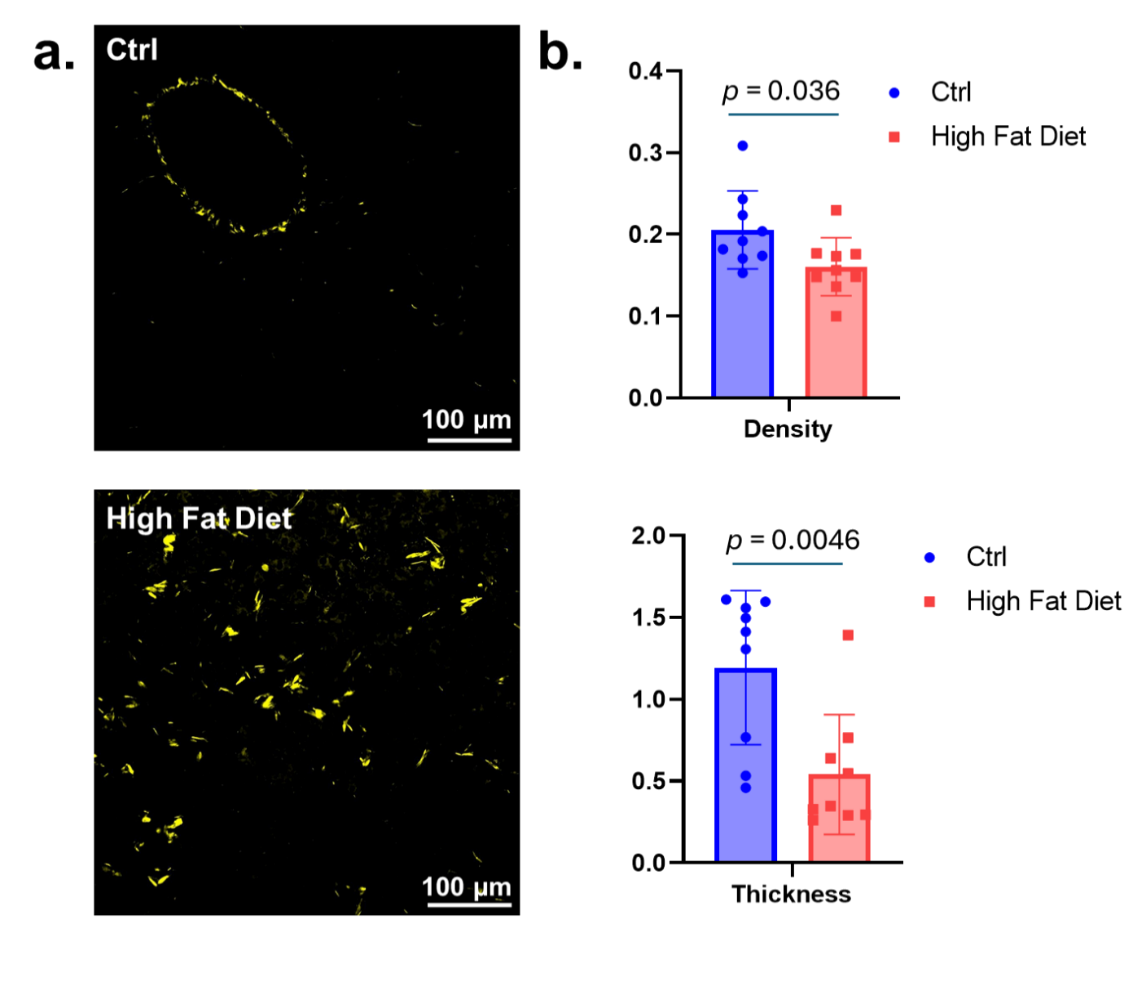

**Fig S4. Liver collagen analysis. a.** SHG images revealed distinct spatial distributions of collagen fibers in liver tissues. In healthy liver tissues, collagen fibers were predominantly localized in the periportal regions near the veins, whereas in high-fat diet liver tissues, collagen fibers were distributed more diffusely throughout the tissue. **b.** Collagen fibers exhibited difference in both density and thickness depending on their spatial context. Collagen fibers in close proximity to hepatocytes display lower density and thinner morphology compared to those associated with the perivascular regions in healthy liver tissues. (n = 9, 3 different samples per each group, 3 ROIs per each sample)

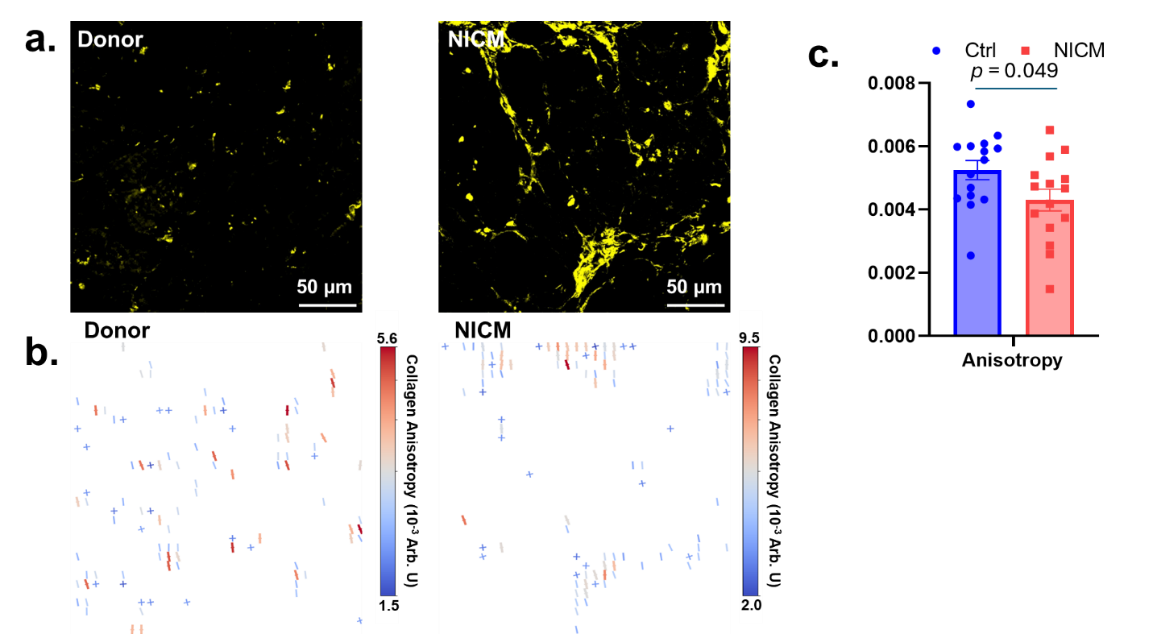

**Fig S5. Collagen analysis in cardiac tissues. a.** The spatial distribution of collagen fibers differs markedly between healthy and NICM samples. In NICM samples, collagen fibers were distributed more extensively in the regions surrounding myocytes. **b.** In addition to altered spatial distribution, collagen fibers in NICM samples exhibited a higher degree of structural alignment compared to those in healthy cardiac tissues. **c.** The increased fiber alignment in NICM samples was clearly visualized in the anisotropy map, where NICM-derived collagen displays lower anisotropy values relative to collagen in healthy tissues, indicating a more ordered and directionally uniform fiber architecture. (n = 15, 3 different samples per each group, 5 ROIs per each sample)

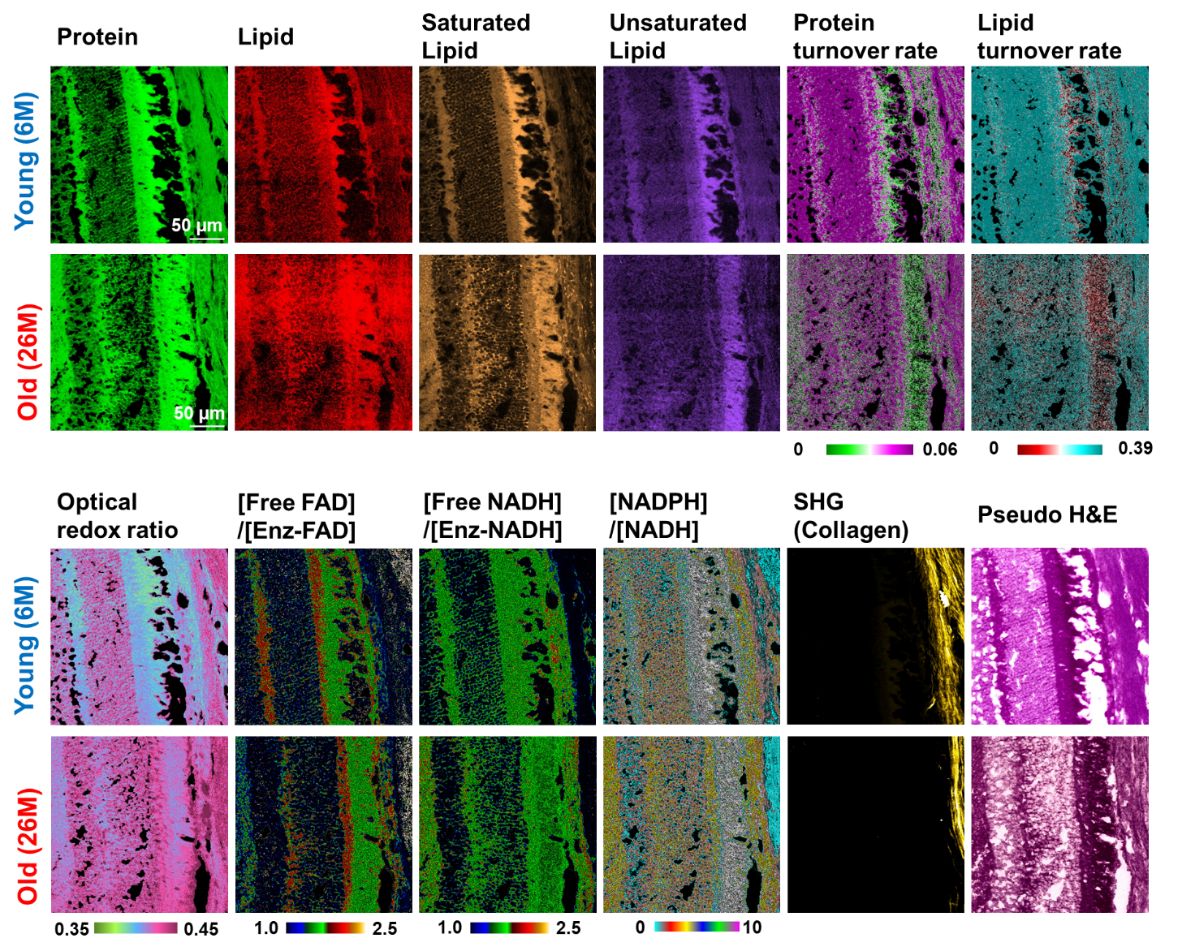

**Fig. S6. Multimodal image channels.** The image channels utilized for retina aging study are listed. As depicted in this figure, multiple image channels representing molecular distribution and metabolic activities are generated from the same ROIs using MANIFEST. Additionally, depending on the analysis pipeline, more channels related to various lipid subtypes or other functional group information can be added.

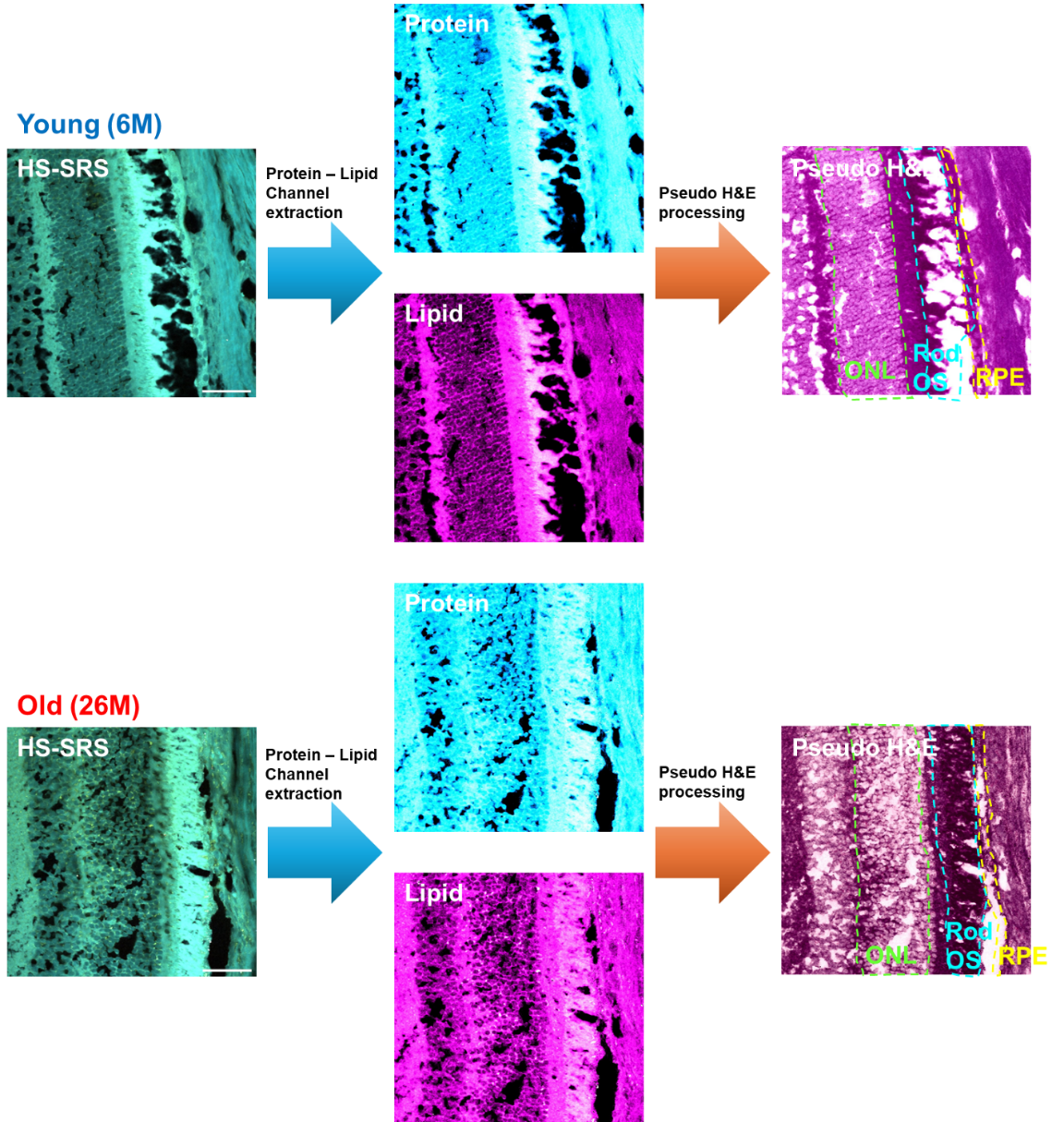

**Fig S7. Pseudo H&E image processing from HS-SRS image stack.** From the HS-SRS image stacks, protein and lipid channel images were extracted by linear spectral unmixing method. The unmixed protein and lipid channel images were used to generate pseudo H&E images.
